# Making sense of chaos: replication and segregation defects in the yeast *mcm4^Chaos^*^3^ mutant

**DOI:** 10.64898/2026.10.30.763219

**Authors:** Jocelyn I. Verhey, Amy K. Moore, Joseph O. Armstrong, Gina M. Alvino, Maitreya J. Dunham, Bonita J. Brewer, M. K. Raghuraman

## Abstract

Genome instability is a defining feature of cancer and is strongly influenced by alterations in the fidelity and timing of DNA replication. The conserved Mini-Chromosome Maintenance (MCM 2-7) helicase is a central component of the eukaryotic DNA replication machinery, playing critical roles in origin licensing and activation. Genetic screens for genome instability mutations in mice identified *mcm4^Chaos3^*, a hypomorphic allele of *MCM4* associated with the formation of micronuclei and tumorigenesis, suggesting that the *mcm4^Chaos3^*allele causes genome instability. To define the mechanistic basis of this instability, we examined the effects of the corresponding mutation in *Saccharomyces cerevisiae.* We found that *mcm4^Chaos3^* selectively compromises origin firing at a subset of early S-phase origins, with pronounced effects at many centromere-proximal origins. Chromosome V, containing the sensitive centromere-adjacent *ARS510,* exhibited elevated loss rates that were rescued by replacement with the robust early-firing origin *ARS305*, restoring both origin activity and chromosome stability. Together these findings demonstrate that *mcm4^Chaos3^* compromises genome stability by impairing a subset of early, centromere-proximal origins that are intrinsically sensitive to origin activation capacity. Our results establish that robust origin function near centromeres is critical for accurate chromosome segregation and reveal how subtle perturbations in helicase function can lead to chromosome instability.

**Author Summary:** The defining feature of cell division is the distribution of replicated chromosomes to the two daughter cells. Each chromosome contains a single centromere that is essential for their faithful inheritance. But before a cell divides, it must duplicate its DNA. DNA replication begins at sites called origins that differ in their activity and regulation. We studied a mutant version of Mcm4, a component of the molecular machinery that unwinds DNA during replication, to determine how compromised replication affects chromosome inheritance. This mutation was first discovered in mice, where it causes chromosome instability and cancer. Using Baker’s yeast, we found that the mutation reduces activity at a subset of origins, with origins near centromeres being preferentially affected. We also found an approximately three-fold increases in the rate that a test chromosome (chr V) is lost. These results suggest that the *mcm4^Chaos3^* mutation can promote genome instability by compromising timely replication of centromeres. Replacing a sensitive origin near the centromere with a robust origin from elsewhere in the genome restored local replication activity and rescued chromosome loss to normal levels. This rescue depended on origin-associated binding sites and a centromere-associated protein, indicating that both origin features and chromosomal context contribute to replication activity. Our findings show that selective defects in replication initiation can compromise chromosome inheritance and that restoring activity at a single centromere-proximal origin can suppress this instability.

## Introduction

The initiation of DNA replication is a tightly regulated process that is essential for the maintenance of genome stability. Control starts at the sequence level where replication initiates at hundreds of discrete sites (origins) along eukaryotic chromosomes which vary in their efficiency and time of initiation during S-phase. The temporal and spatial regulation of origins ensures that the genome is replicated once and only once per cell cycle (1–3). This regulation is achieved through a step wise mechanism: (1) origin licensing in late M/G1 phase, during which the Origin Recognition Complex (ORC), Cdc6 and Cdt1 load the replicative helicase onto origin DNA, and (2) origin firing in S-phase when two kinases, Dbf4-dependent kinase (DDK) and S-phase cyclin-dependent kinase (CDK), activate the helicase to initiate DNA unwinding (4, 5).

The replicative helicase plays a vital role in chromosome maintenance through its control over origin activation. In yeast this helicase is composed of Cdc45, the MCM complex, and the GINS complex, all of which have direct orthologs in metazoans (6). This high conservation across eukaryotes makes *S. cerevisiae* an ideal model for dissecting the structure, function, and regulation of this essential helicase. Helicase activation in S-phase requires DDK-mediated phosphorylation of MCM subunits, which promotes the recruitment of Cdc45 and the GINS complex to form the active CMG (Cdc45-Mcm2-7-GINS) helicase (7). DDK is therefore a rate-limiting factor for origin firing, and its recruitment to individual origins is a critical determinant of origin activation efficiency and timing (8, 9). Because DNA replication is an essential cellular function, it was long considered that mutations in replication initiation proteins would not be tolerated in metazoans. However, studies over the last two decades have revealed that mutations in these proteins have been found in mammals including humans, with diverse consequences. For example, mutations in the pre-Replication Complex (pre-RC) have been identified to cause Meier-Gorlin syndrome. Mutations in helicase proteins have been shown to cause various cancers, so much so that MCM proteins have emerged as effective biomarkers for early detection of many common cancers (10–14).

The MCM2-7 complex is a heterohexameric ring that functions as the core catalytic motor of the helicase (15, 16). The *Chaos3* allele of Mcm4 was originally identified in mice as a dominant, viable mutation that causes chromosome instability and promotes tumorigenesis (11). Introduction of the analogous mutation into budding yeast produces an origin-specific minichromosome maintenance defect: Shima et al. found that plasmids containing the early-firing origin *ARS1* were lost at higher rates in *mcm4^Chaos3^* than plasmids containing the late-firing origin *ARS120*, suggesting that this Mcm4 defect disrupts replication initiation at some origins but not others (11). However, the genomic scope of this differential origin sensitivity has not been characterized. For example, it was unknown whether plasmid maintenance is particularly susceptible to the effect of the *mcm4^Chaos3^*mutation or if chromosomal replication is also affected. Here we demonstrate that chromosomal origins are also compromised and ask whether there is a pattern to which origins are affected and whether the effects on chromosomal origins cause an increase in chromosome instability.

A key step in resolving these questions lies in understanding how DDK is recruited to different origins. While DDK is essential for helicase activation at all origins, the mechanisms by which DDK is recruited to individual origins vary. Centromere-proximal origins, which fire early in S-phase, utilize a specialized pathway mediated by Ctf19, a component of the Ctf19 kinetochore complex (also called COMA) (17, 18). Natsume et al. showed that Ctf19 recruits DDK to the kinetochore during telophase, which in turn promotes the early recruitment of Sld3-Sld7 and the timely activation of flanking origins within approximately 20 kb of the centromere (17). This kinetochore-dependent DDK recruitment pathway explains the previously observed CEN-dependent advancement of origin firing time (19). Furthermore, it ensures that centromere-proximal origins fire robustly and early, coordinating centromere DNA replication with kinetochore function (19). Origins outside this 20 kb window rely on other distinct DDK recruitment mechanisms for helicase activation. The Forkhead (Fkh) transcription factors Fkh1 and Fkh2 bind to Forkhead Binding Sites (FBS) located near the ARS consensus sequence (ACS) of a subset of origins and these Fkh proteins facilitate the recruitment of DDK and promote early origin firing (20, 21). The orientation and spacing of the FBS are critical for Fkh protein recruitment, and disruption of these sites diminishes origin activity (22). The origin *ARS305* is a well-characterized example of a non-centromere-proximal, FBS-dependent early origin, whereas *ARS510* lies near *CEN5* and is dependent on Ctf19 for its early activation (23). The existence of distinct DDK recruitments pathways raises the possibility that the differential sensitivity of origins to *mcm4^Chaos3^* observed by Shima et al. (11) reflects differences in how each origin recruits DDK and activates the MCM helicase.

Early replication of centromeric DNA is itself critical for chromosome stability (24). During S-phase, centromere replication transiently disrupts kinetochore-microtubule attachments, and timely reestablishment of those attachments is important for faithful chromosome segregation (25). Disruption of the normal timing of centromere replication—due to delayed or reduced origin firing—can delay the window during which the kinetochore is detached, increasing the risk of chromosome mis-segregation and aneuploidy (24). Thus, defects in centromere-proximal origin firing may have consequences not only for DNA replication but also for the fidelity of chromosome inheritance.

In this study, we set out to define the broader genomic pattern of origin sensitivity associated with compromised Mcm4 function. We further asked whether differential sensitivity to *mcm4^Chaos3^* correlates with origin location, firing timing, or the pathway through which an origin recruits DDK, with particular attention to Ctf19-dependent centromere-proximal origins and Fkh-dependent early origins. We find that chromosomal origins are sensitive to the *mcm4^Chaos3^* mutation in a similar pattern to those identified by Shima et al. (11). Half of the origins within 10 kb of the centromere are sensitive to the *mcm4^Chaos3^* mutation, and this sensitivity results in an increase in genomic instability, characterized by an increase in Chromosome V loss. Replacing a sensitive, centromere-proximal origin with a robust, non-centromere-proximal origin restores origin function and rescues chromosome instability. Our results reveal that origins are intrinsically sensitivity or robust to the *mcm4^Chaos3^* mutation regardless of location on the chromosome which could be due to differing DDK recruitment mechanisms.

## Results

### The effect of the *mcm4^Chaos3^* mutation on origin firing is not limited to plasmid origins

We first examined general effects of the *mcm4^Chaos3^*mutation by assessing growth rates and by flow cytometry, which provide useful baseline assessments of yeast strain fitness, cell-cycle progression and replication-associated stress. We found that mutant populations grew slightly slower than their wild-type counterparts (wild type: 100.8 & 105.4 min, *mcm4^Chaos3^*: 105.7 & 109.6 min; Sup Fig 1A) and in co-culture wild type cells outcompeted the *mcm4^Chaos3^* mutant (Sup Fig 1B). Flow-cytometric analysis also revealed a subtle delay in S-phase progression in *mcm4^Chaos^* cells, which was largely resolved by 90 minutes (Sup Fig 1C). Primary observations by Shima et al. (11) and the subtle defects of *mcm4^Chaos3^* cells that we discovered through growth assays prompted us to investigate origin activity of *mcm4^Chaos3^* cells and its effect on DNA replication fidelity.

**Figure 1:**
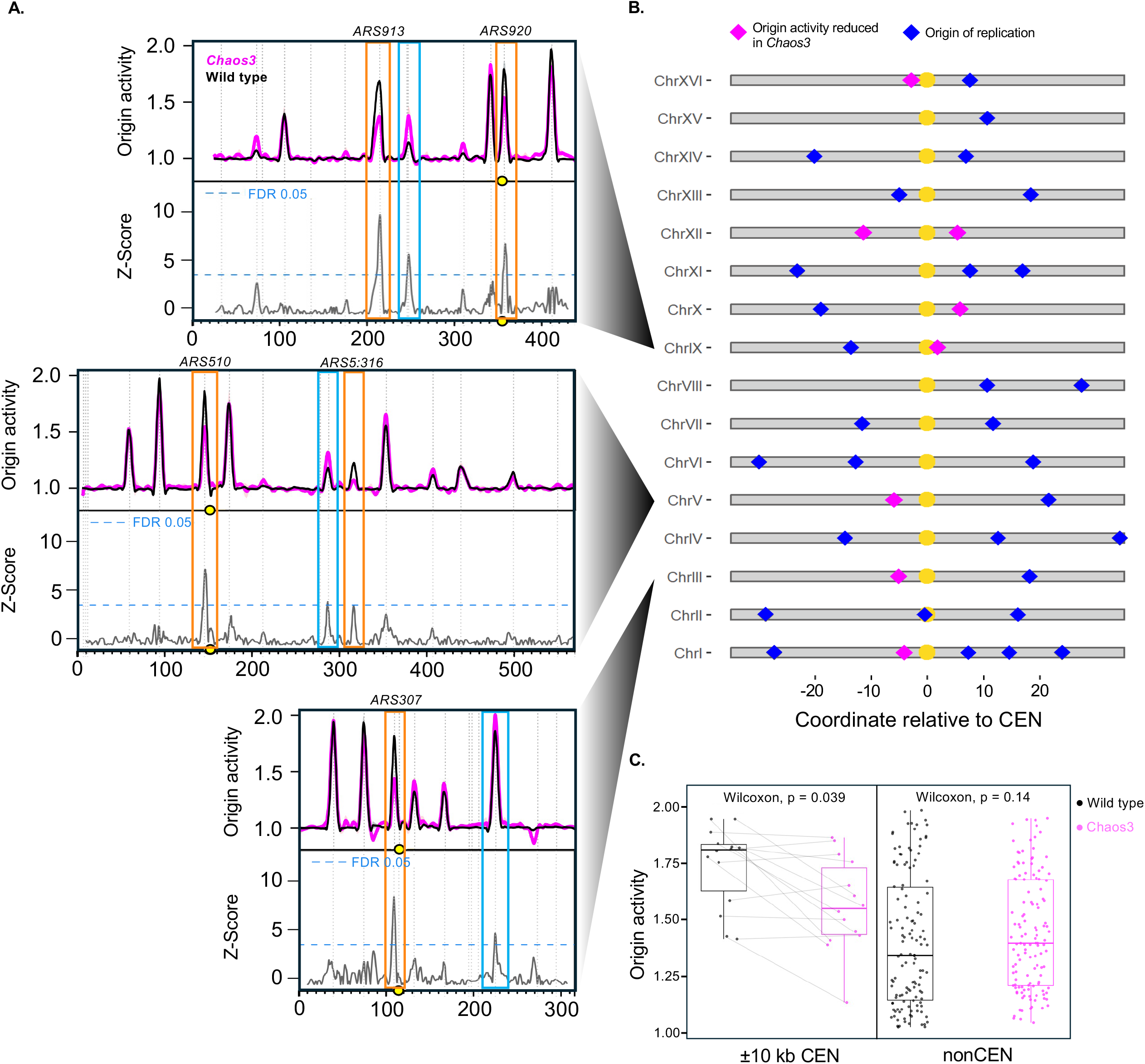
Replication profiles measuring early S-phase origin activity. (A) Replication profiles for WT (black) and *mcm4^chaos3^*(fuchsia) for Chr III, V, and IX. Profiles were generated by measuring the ratio of ssDNA in S phase relative to ssDNA in G1. The x-axis represents chromosomal position (kb), and the centromere is indicated by a yellow circle. Vertical dashed lines indicate locations of confirmed origins from OriDB (62). Orange boxes highlight origins that show decreased activity in the *mcm4^Chaos3^* mutant and blue boxes highlight origins with increased activity in the *mcm4^chaos3^* mutant determined by normalizing the data to Z-score and picking peaks with a FDR < 0.05. (B) Alignment of the centromeres (yellow circle; ± 35 kb of flanking unique sequence) for each of the 16 chromosomes in the genome. Blue diamonds indicate all known origins located within 35 kb on either side of the centromere. Fuchsia diamonds represent origins that have decreased activity as identified in (A). (C) Comparison of origin activity of WT (black dots) and *mcm4^chaos3^* (fuchsia dots) in centromere-proximal (left) and non-centromere-proximal regions(right). Dark gray lines connect the origin activity in wild type to the origin activity in *mcm4^Chaos3^* for the centromere-proximal origins. Origins within 10 kb of the centromere (CEN) show a significant reduction in activity in *mcm4^chaos3^* compared to WT (Wilcoxon test, *p* = 0.039). In contrast, origins located outside the 10 kb CEN window (nonCEN) show no significant difference in activity between genotypes (Wilcoxon test, *p* = 0.14).

Because the MCM complex is crucial for origin initiation as well as for fork elongation, in principle the *mcm4^Chaos3^* mutation could affect either process. However, given Shima et al.’s observation of origin dependent variation in plasmid loss rates (11), we hypothesized that at least in yeast, the primary effect of the mutation would be on origin firing. To determine whether the effect of the *mcm4^Chaos3^* mutation extends to chromosomal origins, we used an assay that measures early firing chromosomal origins in *S. cerevisiae* (*26, 27*). This assay relies on the observation that hydroxyurea (HU) limits the expansion of the nucleotide pool (28, 29) so that cells grown in such conditions exhibit slowed replication fork progression, exposing unreplicated, single-stranded DNA (ssDNA) immediately adjacent to active replication origins (30). We labeled these regions of single-stranded DNA with fluorescent dyes and hybridized the labeled DNA to a microarray to generate origin activity profiles. In such profiles, the x-axis corresponds to position along a representative chromosome and the y-axis to origin activity, with peaks indicating active, early S-phase origin firing; the landscape of peak heights thus reflects differences in origin activity, with the tallest peaks marking the most active origins. Overlaying replication profiles allows comparison of origin activity across genotypes and mutational backgrounds. We performed two replicates of the experiment. Comparison of wild-type and *mcm4^Chaos3^*profiles revealed a variety of affected origins (Figure 1A, Sup Fig 2). Even by manual inspection, we see that most early firing origins are unaffected by the presence of the *mcm4^Chaos3^* mutation. However, of the origins that are affected we see both increases (Figure 1A, blue boxes) and decreases in origin activity (Figure 1A, orange boxes). To assess these differences systematically, we converted the absolute difference between wild type and mutant profiles to Z-score (units of standard deviation) by normalizing them against same-strain differences and estimating a Z-score cutoff for a false discovery rate of 0.05 (see Materials and Methods) (Figure 1A, Sup Fig 2). Z-scores allowed us to identify those origins that were significantly altered in their replication efficiency in the *mcm4^Chaos3^*mutant. Most notably, we see a significant reduction in origin activity at ∼20 of the ∼150 early S-phase origins (Figure 1A, orange boxes).

**Figure 2:**
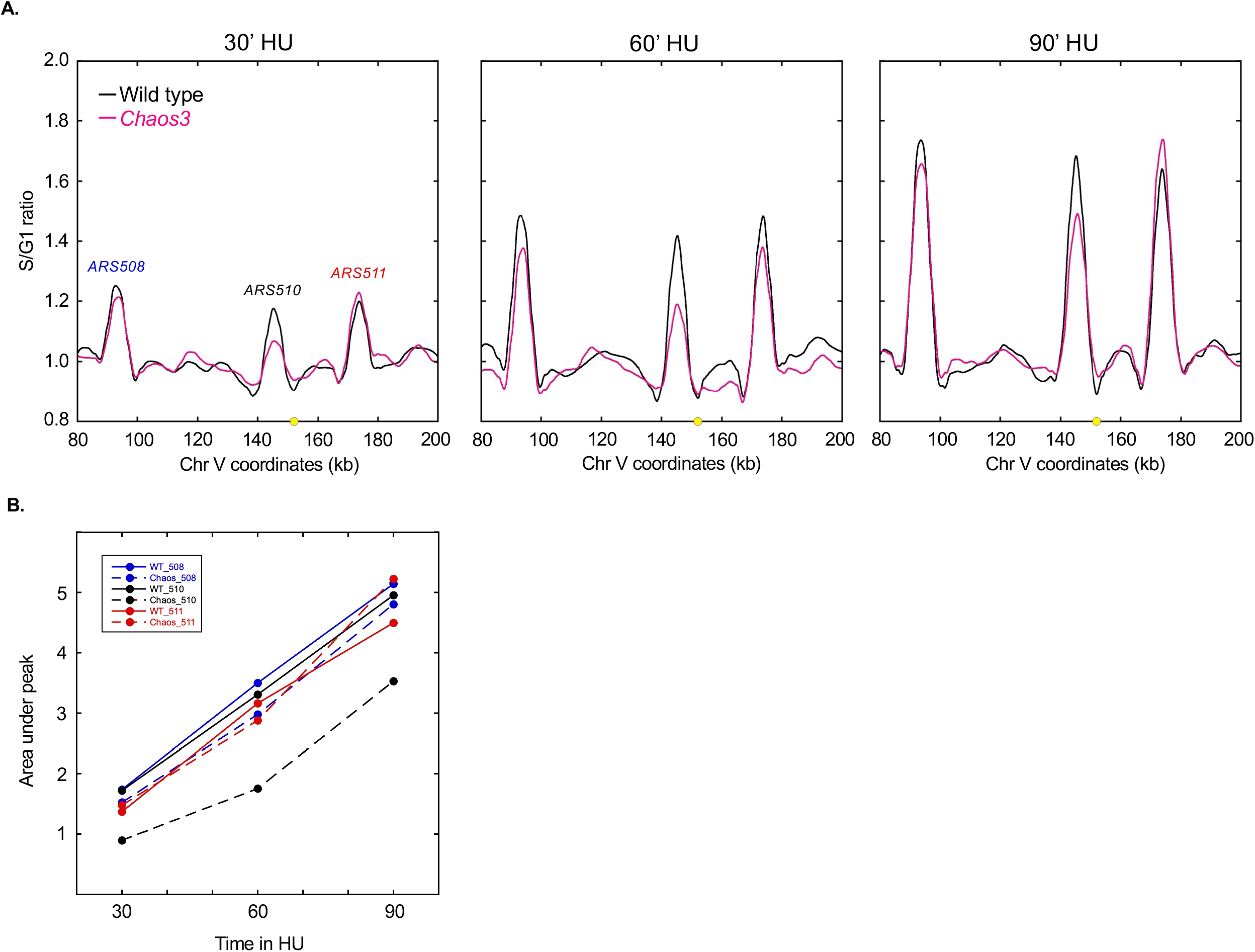
Replication initiation defect of *mcm4^Chaos3^* sensitive origin. (A) Comparison of short read sequencing profiles of replication progression from a hydroxyurea (HU) synchronized 30-, 60-, 90-minute time course for wild type (black line) and *mcm4^Chaos3^* (fuchsia line) for a representative portion of chromosome V (x-axis coordinates). The centromere is indicated by a yellow circle, and S-phase relative to G1 read depth is plotted on the y-axis. (B) Origin replication activity plot of the HU time course for *ARS508* (blue), *ARS510* (black), and *ARS511* (red) as indicated in (A). Area under the peak values (y-axis) for 30, 60, and 90-minute samples (x-axis) are plotted as colored circles with either solid (WT) or dotted lines (*mcm4^Chaos3^*).

### Centromere-proximal origins are preferentially affected by the *mcm4^Chaos3^* mutation

When we examined the location of these ∼20 *mcm4^Chaos3^*affected origins in the genome we noticed a non-random clustering of origins that mapped within ∼10 kb of the centromere (Figure 1B). We generated a map of origin activity for each of the 16 chromosomes in the genome, plotting origins on a coordinate system centered on each centromere and narrowing the view to a 70 kb window (± 35 kb) around the centromere (Figure 1B). We find that half of the origins within ∼10 kb of the centromere (centromere-proximal) show reduced origin activity in the *mcm4^Chaos3^* background. Compared to wild-type, centromere-proximal origins exhibit a statistically significant reduction in activity in *mcm4^Chaos3^* (Wilcoxon test, p = 0.039), whereas non-centromere-proximal origins show no significant difference (p = 0.14) (Figure 1C).

To further confirm the reduction in centromere-proximal origin activity, we performed short-read sequencing across 30-, 60-, and 90-minute timed samples of cells released from a G1 arrest into S phase in HU. When comparing wild-type and *mcm4^Chaos3^*HU read-depth profiles we found a slight reduction in replication after 60’ in HU for the three highly active origins surrounding *CEN5* on chromosome V (Figure 2A). The two flanking origins recovered and reached the same level of origin activation as wild type, or better, by 90’ but *ARS510*, the centromere-proximal origin identified in Figure 1A, failed to recover to wild type levels of origin firing in the same time period. Plotting the area under the peak for the representative origins for each time point recapitulated the delay in replication at 60’ and highlights *ARS510* failure to recover to wild type replication levels as seen in the read-depth profiles (Figure 2B). This result suggests that while most origins may suffer a slight delay in origin firing in the *mcm4^Chaos3^* mutant, *ARS510* and other centromere-proximal origins suffer an overall decrease in early origin firing (Figure 2B, Sup Fig 3). The reduced initiation at early centromeric origins could result in a delay in centromere replication in a proportion of *mcm4^Chaos3^* cells, as previously shown for *ctf19Δ* and a strain containing a tagged version of Dbf4 with *myc* at its C-terminus (23, 31).

**Figure 3:**
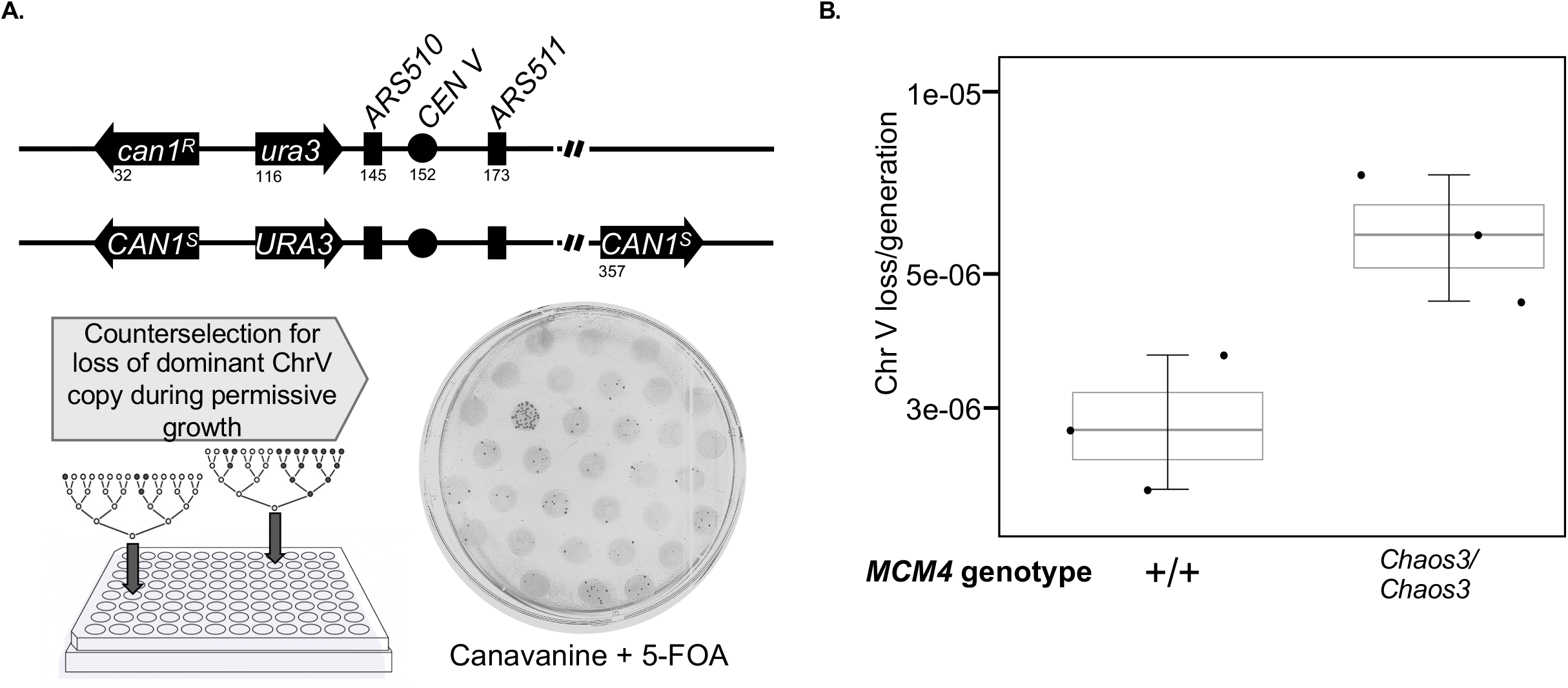
Whole chromosome loss due to delay in centromere replication. (A) Illustration of the heterozygous Chr V used to measure whole chromosome loss rates in various mutational backgrounds. The top “recessively marked” copy of Chr V contains *can1^r^* (conferring canavanine resistance) and *ura3* (conferring uracil auxotrophy). The bottom “dominantly marked” copy of Chr V contains *CAN1^S^* (conferring sensitivity to canavanine) and URA3 (conferring uracil prototrophy). The dominantly marked copy was initially selected for by growth in media lacking in uracil. Nonselective growth in synthetic complete media allowing for stochastic loss of the dominantly marked (bottom) copy before counterselection on canavanine + 5-FOA 15 cm plates to identify cells that had lost the dominant copy. (B) Whole Chromosome V loss rate measured in diploids using heterozygous *CAN1^S^/can1^r^* and *URA3/ura3* as depicted in (A), via Luria-Delbrük fluctuation assay. Loss rates for each strain were estimated using three biological replicates, with each estimate represented by a dot within the boxplot spanning the interquartile range. The *MCM4* genotype for the representative diploids is either homozygous *MCM4* (+/+) or homozygous *mcm4^chaos3^* (*Chaos3/Chaos3*). Diploids homozygous for *mcm4^chaos3^* exhibit approximately 3-fold greater Chr V loss compared to WT.

### The *mcm4^Chaos3^* mutation increases the rate of chromosome loss

During centromere replication the kinetochore must entirely detach from the centromeric DNA and subsequently reassemble to facilitate spindle microtubule attachment (25). A disruption in the timing of kinetochore reattachment due to delayed centromere replication exposes chromosomes to an increased risk of chromosome mis-segregation and aneuploidy (24). Thus, if the timely, robust replication of the centromere is essential for proper chromosome segregation then the subset of cells that are unable to initiate replication from centromere-proximal origins such as *ARS510* might be compromised in segregation of chromosomes such as Chr V during mitosis. To test for increased chromosome loss, we turned to analysis of homozygous *mcm4^Chaos3^*diploids.

Our chromosome loss assay is based on the differential sensitivity of the two copies of chromosome V to canavanine. To minimize false positives arising from post-plating mutations (32, 33), we first constructed a “dominantly marked” test chromosome V by inserting an additional *CAN1^S^*marker on the arm opposite the native *CAN1^S^* and *URA3* markers (see Materials & Methods). By inserting the additional copy of *CAN1^S^* on the arm opposite the native copy we were able to detect instances of whole chromosome loss rather than chromosome fragmentation. This strain was mated to yeast carrying a “recessively marked” chromosome V homolog marked with *can1^R^* and *ura3*. The heterozygous *CAN1* and *URA3* diploid enabled chromosome loss to be tracked by counterselection (Figure 3A, chromosome diagram). To detect chromosome V loss, cells were first grown to saturation under conditions that maintained the dominantly marked chromosome V, then diluted into permissive medium to allow chromosome loss events to occur. Each culture was then plated in its entirety on counter-selective medium containing canavanine and 5-FOA, on which only cells that had lost the dominantly marked chromosome V copy and retained the recessively marked homolog could grow (Figure 3A). Examining three biological replicates for each genotype we found that diploids homozygous for the *mcm4^Chaos3^* mutation exhibited an approximately three-fold increase in chromosome V loss per generation compared with wild-type cells (Figure 3B). The increase in chromosome loss reflects how the delay in centromere replication due to the decrease in *ARS510* activity affects the fidelity of chromosome maintenance, underscoring the importance of robust centromere-proximal origin activity.

### Robust origin replacement rescues origin firing and chromosome loss rates

The centromere has been shown to affect the temporal landscape of origin firing (19) with proximity to the centromere directly affecting origin activity and timing. Pohl et al.’s, ectopic centromere experiment (19) demonstrated that a late replicating region of a chromosome becomes early replicating based solely on the relocation of a functional centromere to the late replicating region, establishing that the presence of a functional centromere is sufficient for early activation of adjacent origins. This example of the centromere’s influence on origin activity led us to ask if the particular sensitivity of centromere-proximal origins to the *mcm4^Chaos3^*mutation was due to proximity to the centromere itself—i.e., whether any origin placed at that location would be compromised in the *mcm4^Chaos3^*background. To test this hypothesis, we replaced the *mcm4^Chaos3^* sensitive origin *ARS510* with the robust, non-centromere-proximal *ARS305*. This replacement was completed by substituting a 421-bp region encompassing *ARS510* on chromosome V with a 653-bp fragment containing *ARS305* and its flanking sequence, generating the *ars510Δ::ARS305* allele (Figure 4A, chromosome diagram).

**Figure 4:**
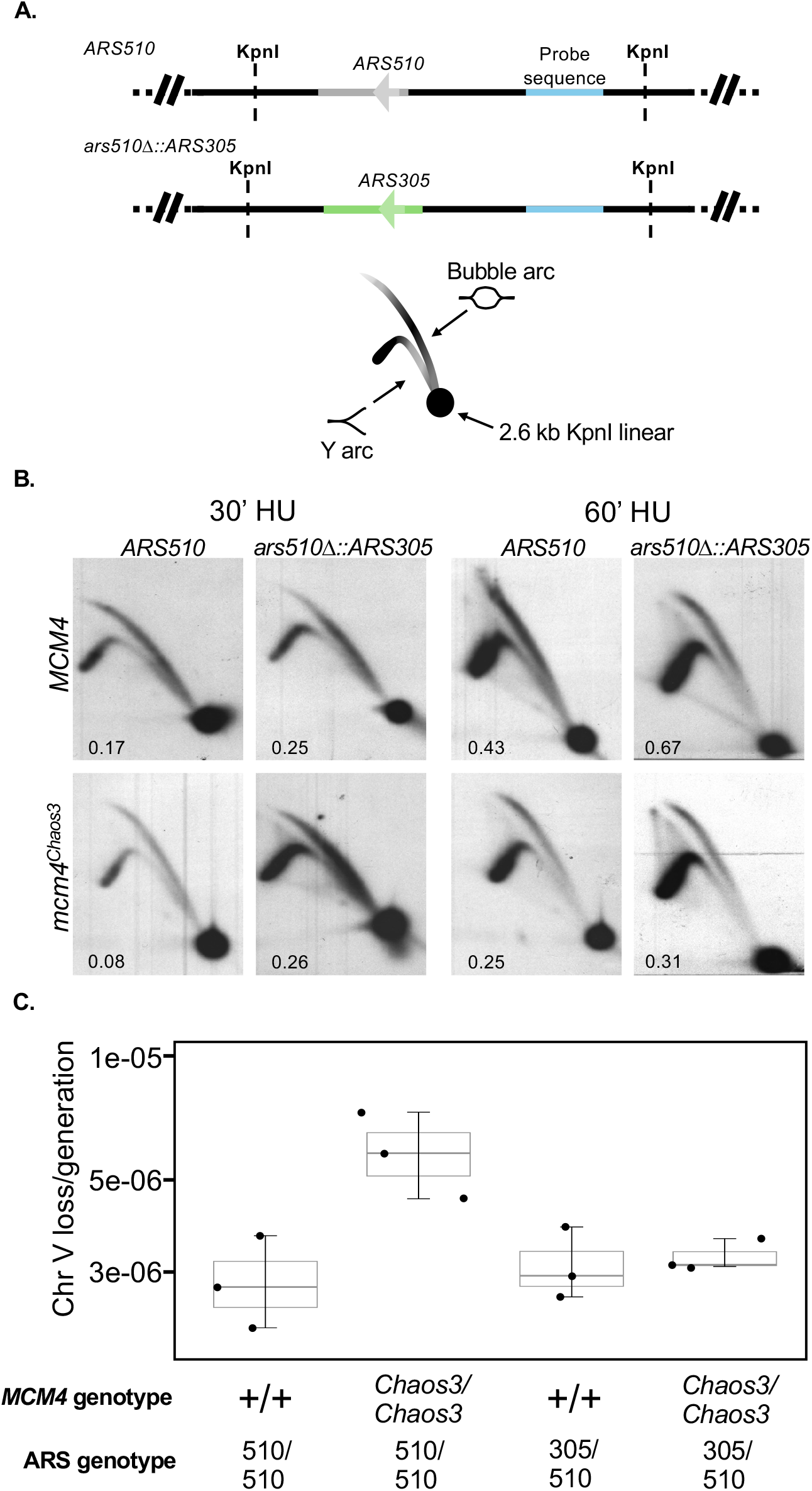
Origin activity and chromosome loss rates for an origin replacement mutant. (A) Diagrams of the region on Chr V containing the *mcm4^chaos3^*sensitive origin, *ARS510,* and the replacement construct *ars510Δ::ARS305*, in which ARS*510* is replaced by the robust non-centromeric origin *ARS30*5. The arrow depicts the orientation of the T-rich strand of the ACS; directionality was maintained in the origin replacement. The restriction enzyme KpnI cuts this region of Chr V twice creating a ∼2.6 kb fragment used to probe (sky blue “probe sequence”) origin activity at the *ARS510* locus by 2D gel electrophoresis. As replication initiates, replication bubbles extend from the origin and collect in the upper bubble arc of the 2D gel diagram. Because the origin is not centered in the digestion fragment, replication bubbles can extend past either KpnI site and create forked intermediates that collect in the lower Y arc. Any non-replicating DNA remains linear and collects in the 1N spot. The color gradient in the 2D gel diagram represents the progression of replication intermediates. (B) 2D gels analysis of *MCM4* and *mcm4^chaos3^*after 30 minutes or 60 minutes of growth in hydroxyurea (HU), comparing *ARS510* and *ars510Δ::ARS305*. At 30 minutes in HU, ARS510 origin activity in the *mcm4^chaos3^* background is approximately half that of wild type (0.08 cs 0.17). Replacement of *ARS510* with *ARS305* (*ars510Δ::ARS305*) restores origin activity in the *Chaos3* double mutant (30’ HU 0.25 vs 0.26). (C) Luria-Delbrük fluctuation assay to measure whole chromosome V loss rates as described in Fig. 2B. The right two box plots show the heterozygous *ars510Δ::ARS305/ARS510* Chromosome V has comparable levels of loss in homozygous *MCM4* and homozygous *mcm4^chaos3^* strains as well as the baseline WT levels of loss observed in the left most box plot (+/+, 510/510).

We examined origin activity at the replacement locus in *MCM4* and *mcm4^Chaos3^* cells by two-dimensional (2D) gel electrophoresis. 2D agarose gels of digested genomic DNA from cells released from G1 into HU were Southern blotted and probed using the specified region on the digestion fragment as indicated in Figure 4A. Replication intermediates produced by origin firing migrate as bubbles and collect in the top arc, bubbles that have extended past one of the digestion sites migrate as forked intermediates and collect in the lower Y arc and non-replicating linear DNAs define the 1N spot (Figure 4A). Because HU limits replication fork progression, all replication intermediates (bubbles and Ys) detected at this locus reflect local origin firing rather than passive replication from neighboring origins. The signal in the two arcs is combined and divided by the signal produced in the 1N spot to determine overall activity at this locus. 2D gel comparison for *ARS510* of *MCM4* and *mcm4^Chaos3^*backgrounds recapitulate the data seen in the ssDNA replication profiles: the activity of *ARS510* is reduced by half in *mcm4^Chaos3^* compared to wild type (0.08 vs 0.17 at 30’ HU; 0.25 vs 0.43 at 60’ HU) (Figure 4B). Replacement of *ARS510* with *ARS305* successfully restored origin activity to wild-type levels at that location, with the replacement strain showing comparable activity in both *MCM4* and *mcm4^Chaos3^* backgrounds (0.25 and 0.26 at 30’ HU) (Figure 4B). This result suggests that the centromere’s effect on origin sensitivity, if any, is limited to specific origins and that *ARS305* maintains an intrinsic resistance to the *mcm4^Chaos3^* mutation regardless of location in the genome.

To test whether restoring origin firing rescues chromosome loss we repeated the chromosome loss assay with the *ars510*Δ::*ARS305* replacement strain. We find that in the replacement strain, *MCM4* and *mcm4^Chaos3^* have comparable chromosome V loss rates, both of which are comparable to wild type (Figure 4C). The rescue of chromosome V loss in the *ars510*Δ::*ARS305* replacement strain further supports our hypothesis that chromosome stability is directly influenced by the robust, timely replication of the centromere and shows that even subtle perturbations to origin activity can have profound effects on chromosome loss rates.

### Linking the replication defects caused by the *mcm4^Chaos3^* mutation to chromosome mis-segregation using single-origin plasmids

Replication profiles and chromosome loss assays provide a view of the cellular response to the *mcm4^Chaos3^* mutation, but the proximity of neighboring origins could dampen the magnitude of the effect the mutation has on origin activation and genome stability. For example, in the subset of cells where *ARS510* is not active, the flanking origins could compensate for loss of *ARS510* activity potentially masking the effect the decrease in origin activation of *ARS510* has on genome stability. To remove potentially confounding events due to neighboring origins we returned to the plasmid loss assay, as single-origin plasmids allow us to evaluate an origin’s intrinsic sensitivity to the *mcm4^Chaos3^* mutation separate from its native genomic context.

Estimating plasmid loss rates by selective and nonselective growth in combination with selective and nonselective plating is the standard, robust method for estimating plasmid loss; however, this approach cannot differentiate between plasmid loss due to replication errors and loss due to a segregation defect. In other words, plating reveals total plasmid loss rates. Our specific interest in replication initiation led us to use a combination of Southern blotting and plating to parse the contribution of replication versus segregation defects on plasmid loss. We inserted a PCR fragment containing *ARS510* or *ars510Δ::ARS305* into a plasmid containing *CEN5* and the selectable marker *URA3* (p510-CEN or p510.305-CEN, respectively). We transformed wild type and *mcm4^Chaos3^* haploid strains with each of these plasmids and grew the strains in selective and nonselective conditions for ten generations. We plated aliquots of the initial and final cultures on selective and non-selective plates to determine plasmid loss rates and also extracted total DNA from these samples and digested the DNA with a restriction enzyme. We ran the digested DNA on an agarose gel to separate digestion fragments, Southern blotted, and probed with the same probe used for our 2D gel quantification (see Figure 4A). This probe binds to restriction fragments from the region of chromosome V and the plasmid that differ in size, allowing us to quantify both simultaneously. Importantly, this method of quantification is robust to cases of plasmid loss due to segregation defects. Because we collect, digest, and quantify the total amount of DNA present in a representative culture, plasmid copies that were replicated but failed to segregate to a daughter cell are still captured. Thus, the DNA based method specifically reports on loss due to replication failure.

Total loss rates determined from the plating experiment with p510-CEN in the wild type and *mcm4^Chaos3^* backgrounds (Fig 5A, blue bars, left) mirror our findings for Chr V loss rates (Fig 4C): plasmid loss rate is roughly 3-fold higher in the *mcm4^Chaos3^* background. The replacement of the origin in plasmid p510.305-CEN significantly reduces plasmid loss rate in the *mcm4^Chaos3^* mutant (Fig 5A, blue bars, right). This result recapitulates the trend seen in the chromosome loss experiments (Figure 4C).

**Figure 5:**
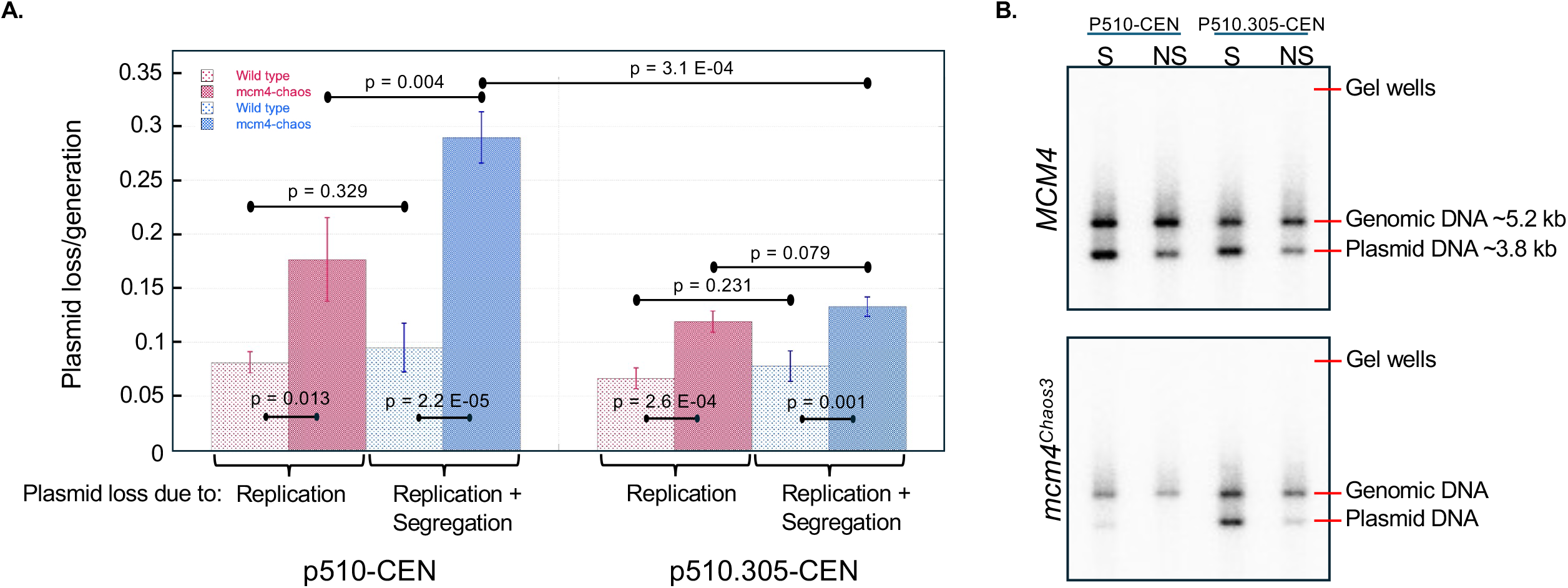
Single-origin plasmid maintenance in wild-type vs *mcm4^Chaos3^* cells. (A) WT and *mcm4^Chaos3^*haploid strains were transformed with a CEN-based plasmid carrying the selectable marker URA3, *CEN5*, and either *ARS510* (p510-CEN) or *ARS305* (p510.305-CEN). DNA was extracted from cultures grown in selective (S) and nonselective (NS) conditions and digested with PstI-HF. When resolved on an agarose gel and probed for the region of Chr V illustrated in Fig 3A two bands are present corresponding to native Chr V (genomic DNA, ∼5.2 kb) and to the plasmid used (plasmid DNA, ∼3.8 kb). The ratio of plasmid to genomic DNA signal under selective versus nonselective conditions is used to quantify plasmid loss attributable to replication errors. (B) *ARS510* and *ars510Δ::ARS305* plasmid loss due to replication and segregation errors (blue boxes) and plasmid loss due to replication errors (pink boxes). Two-tailed unequal variance Welch’s t-test was used to determine statistical significance between various groups as identified by black lines.

Plasmid loss using Southern blots is based on comparing the intensity of the plasmid band to the chromosomal band in the initial and final cell samples (Figure 5B). In the *mcm4^Chaos3^* background, the p510-CEN plasmid band is nearly absent after ∼10 generation of nonselective growth, whereas the p510.305-CEN plasmid band remains visible, consistent with greater loss of the *ARS510*-containing plasmid. Based on quantification of plasmid and genomic DNA bands in selective and non-selective conditions of four biological replicates (Sup Fig 4), we determined mean plasmid loss rates (Figure 5A, pink bars).

The difference between total plasmid loss and replication-based plasmid loss reveals the proportion of loss attributable to segregation-based errors. For p510-CEN, total plasmid loss in *mcm4^Chaos3^* (0.29/generation) substantially exceeded replication-based plasmid loss (0.17/generation), highlighting the significant proportion of plasmid loss due to segregation defects (Figure 5A, left dark pink vs left dark blue bars). In contrast, for p510.305-CEN, replication-based plasmid loss and total plasmid loss were roughly equivalent meaning that loss events due to replication errors can explain the majority of loss events for this plasmid regardless of *MCM4* genotype (Figure 5B, right dark pink vs right dark blue bars). Comparison of plating vs Southern blotting reveal the significant burden segregation defects have on plasmid stability for p510-CEN in the *mcm4^Chaos3^* background. This result emphasizes how some feature of *ARS305* enables the origin to remain a robust origin of replication despite mutational backgrounds and chromatin contexts.

### Centromere replication and chromosome loss in *mcm4^Chaos3^* mutants resembles that of a CEN-origin activation-deficient mutant, *ctf19Δ*

The data thus far point to errors in DNA replication initiation as the cause for the differential origin sensitivity to the *mcm4^Chaos3^*mutation. To investigate the mechanisms that underlie the differences between these origins, we examined pathways by which origins recruit the rate-limiting Dbf4-dependent kinase (DDK) for the phosphorylation of MCM proteins and helicase activation in S-phase (8, 9). Origins within ∼20 kb of the centromere utilize Ctf19, a kinetochore component that recruits DDK to the kinetochore itself, resulting in the activation of the nearest flanking origins (17). To determine whether the mcm*4^Chaos3^*mutation affects the same pathway through which *CTF19* acts, we performed ssDNA profiling on a *ctf19Δ* knockout strain. The *ctf19Δ* mutation was epistatic to *mcm4^Chaos3^*: while we could not observe an additive effect in the *mcm4^Chaos3^ ctf19Δ* double mutant (Figure 6A, purple line), we found that many of the origins with reduced activity in *mcm4^Chaos3^* are reliant on *CTF19* for their activation (Figure 6A, magenta vs orange lines). To assess whether this overlap between *Chaos3*-sensitive and *CTF19*-dependent origins holds true genome wide, we aligned replication profiles from all 16 chromosomes centered at their respective centromere (Figure 6B). Nearly all origins with ± 10-20 kb of the centromere show reduced activity in *ctf19Δ* (orange triangles) and included all of the origins that were sensitive to *mcm4^Chaos3^*(pink diamonds). Origins outside this centromere-proximal window were largely unaffected in either mutant (Figure 6A).

**Figure 6:**
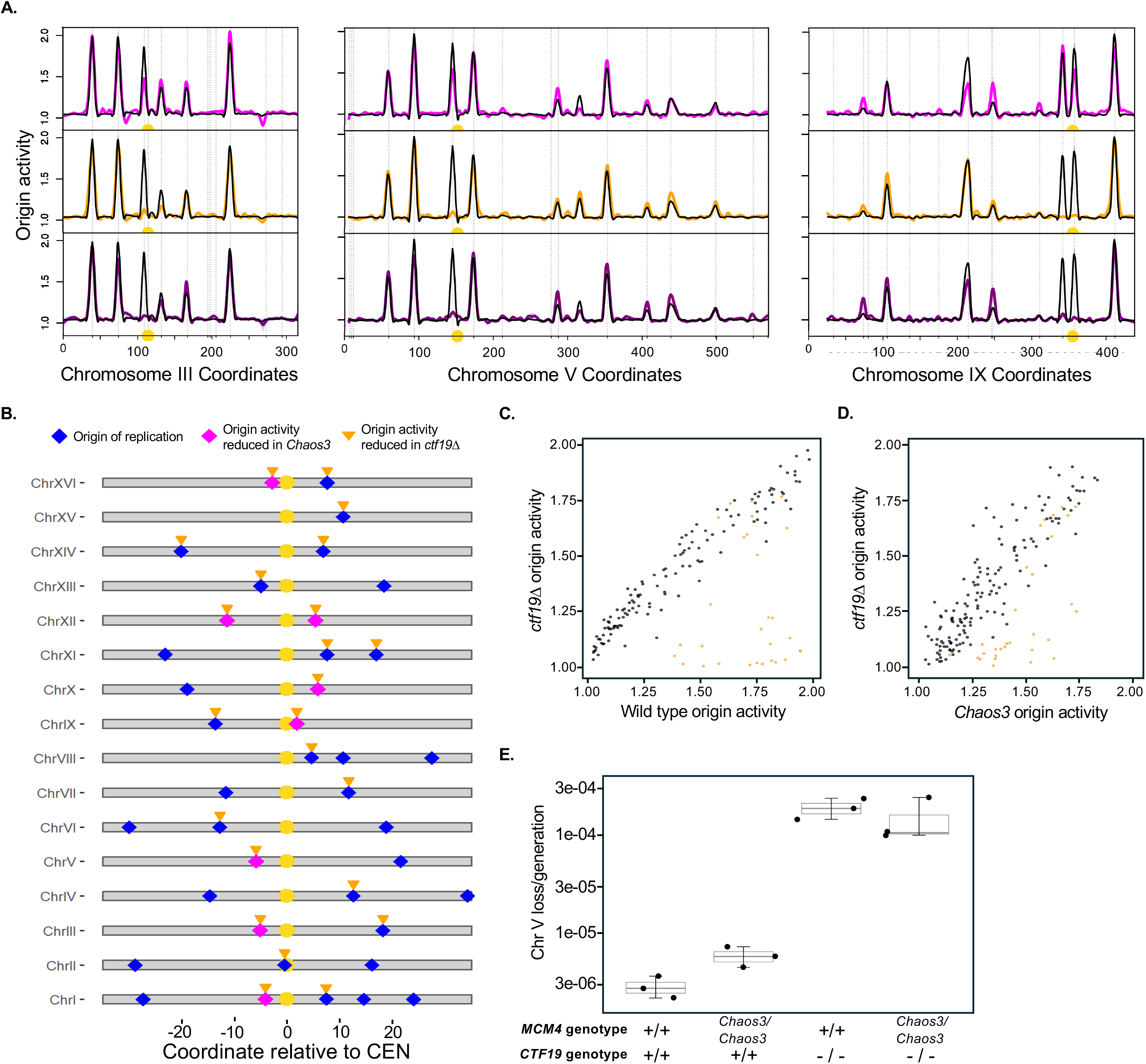
CEN origin sensitivity to replication initiation defect mutation. (A) Replication profiles of Chr III, V, and IX for the wild type strain (black lines) relative to *mcm4^Chaos3^* (fuchsia, reproduced from Fig 1A), *ctf19Δ* (yellow line), and a*ctf19Δ mcm4^Chaos3^* (purple line). The x-axis represents chromosomal position in kb, with the centromere indicated by a yellow circle. Vertical dashed lines indicate locations of confirmed origins from OriDB (62). (B) Diagram of a 70 kb region centered on the centromere (yellow circle) for each of the 16 chromosomes in the genome reproduced from Fig 1B with the addition of origins affected in a *ctf19Δ* mutant (orange triangles). Blue diamonds indicate origins located within 35 kb on either side of the centromere. Fuschia diamonds mark origins with decreased activity as identified in *mcm4^Chaos3^*and are reproduced from Fig 1B. (C) Scatter plot for the relationship between wild-type origin activity (x-axis) and *ctf19Δ* origin activity (y-axis). Black points represent non-centromeric (non-CEN) origins and orange points represent centromeric (CEN) origins. (D) Scatter plot for the relationship between *mcm4^Chaos3^* origin activity (x-axis) and *ctf19Δ* origin activity (y-axis). (E) Box and whisker plots showing the frequency of whole-chromosome V loss in homozygous diploid strains carrying either *MCM4* (+/+) or *mcm4^Chaos3^* (Chaos3/Chaos3) in combination with either *CTF19* (+/+) or *ctf19Δ* (-/-). Individual biological replicates are shown by black points. Boxes represent the interquartile range with the median indicated by the horizontal line, and whiskers represent the data range. The y-axis is plotted on a logarithmic scale to accommodate the broad range of chromosome loss frequencies.

We quantitatively tested the correlation between *CTF19* dependence and *mcm4^Chaos3^* sensitivity across replication origins by comparing origin activity levels across the different genotypes. In a scatter plot of wild-type versus *ctf19Δ* origin activity, non-CEN origins (gray) clustered tightly along the diagonal, indicating no effect of *ctf19Δ*, whereas CEN-proximal origins (orange) deviated markedly downward, reflecting their high activity in wild type but near complete loss in *ctf19Δ* (Figure 6C). When we plotted *Chaos3* origin activity against *ctf19Δ* origin activity, CEN origins again clustered in the lower region, confirming that the same origins compromised by the *mcm4^Chaos3^* are also dependent on Ctf19 (Figure 6D). Non-CEN origins remained largely correlated along the diagonal, consistent with their independence from both mutations.

Having established that *mcm4^Chaos3^* and *ctf19Δ* affect overlapping sets of centromere-proximal origins, we asked whether this shared origin activation defect translates to similar chromosome loss phenotypes. We assayed chromosome V loss rates in wild type, *mcm4^Chaos3^*, *ctf19Δ*, and the *mcm4^Chaos3^ ctf19Δ* double mutant (Figure 6E). As shown above, *mcm4^Chaos3^* caused a modest increase in chromosome V loss relative to wild type. In contrast, *ctf19Δ* caused a dramatic increase in chromosome V loss (∼2 x 10^-4^ per generation), approximately 40-fold above wild type. Other studies of *ctf19Δ* mutants showed similar, but more extreme chromosome loss rates (31, 34, 35). Because we are only investigating the loss rate of chromosome V, but each chromosome has significant defects in centromere-proximal origin firing (Figure 6A), it is likely that the chromosome V loss rate we have determined is significantly lower than the true loss rate due to increased loss of other chromosomes, resulting in cellular lethality prior to detecting loss of chromosome V. The *mcm4^Chaos3^ ctf19Δ* double mutant exhibited a chromosome loss rate (∼1.5 x 10^-4^ per generation) comparable to *ctf19Δ* alone, consistent with the epistasis observed at the level of origin firing. The substantially greater chromosome loss in *ctf19Δ* compared to *mcm4^Chaos3^*, despite their overlapping origin activation defects, suggests that complete loss of *CTF19*-dependent origin activation—or additional *CTF19* kinetochore functions beyond DDK recruitment—contributes to far greater chromosome instability than the partial compromise caused by *mcm4^Chaos3^*.

### Robust nonCEN origin rescues origin activity at an *mcm4^Chaos3^* affected locus and rescues chromosome loss in a Fkh 1/2 binding site dependent manner

As shown previously, most centromere-proximal origins are reliant on Ctf19 for the recruitment of DDK and subsequent phosphorylation and activation (17–19). *ARS305* in its native context is non-centromere-proximal and Ctf19 has no effect on its activation; instead, it utilizes forkhead (Fkh) binding sites (FBS) which recruit Fkh transcription factors for the subsequent recruitment of DDK (20, 21). The FBS were included in the surrounding region for the replacement of *ARS510* with *ARS305*, giving this ectopic origin two methods of DDK recruitment: Ctf19 and FBS (Figure 7A). To determine whether the rescue of the *ARS510* locus by *ARS305* was due to increased DDK recruitment from both pathways or some other feature of *ARS305* itself, we constructed mutant strains utilizing only one method of DDK recruitment. First, we mutated the FBS. The orientation and spacing between the two FBS are essential for the function and recruitment of Fkh proteins (22). As shown in Reinape et al., inversion of the 3’ FBS substantially decreases the recruitment of Fkh proteins (22). We similarly mutated the 3’ FBS in our *ars510Δ::ARS305* replacement strain to disrupt Fkh-dependent DDK recruitment via this pathway (Figure 6A) leaving *ARS305* at its ectopic chromosome V location to rely solely on Ctf19-mediated DDK recruitment. Separately, we combined the *ars510Δ::ARS305* replacement (with FBS still intact) with a *ctf19Δ* knockout, so that only FBS-dependent DDK recruitment remains at this locus.

**Figure 7:**
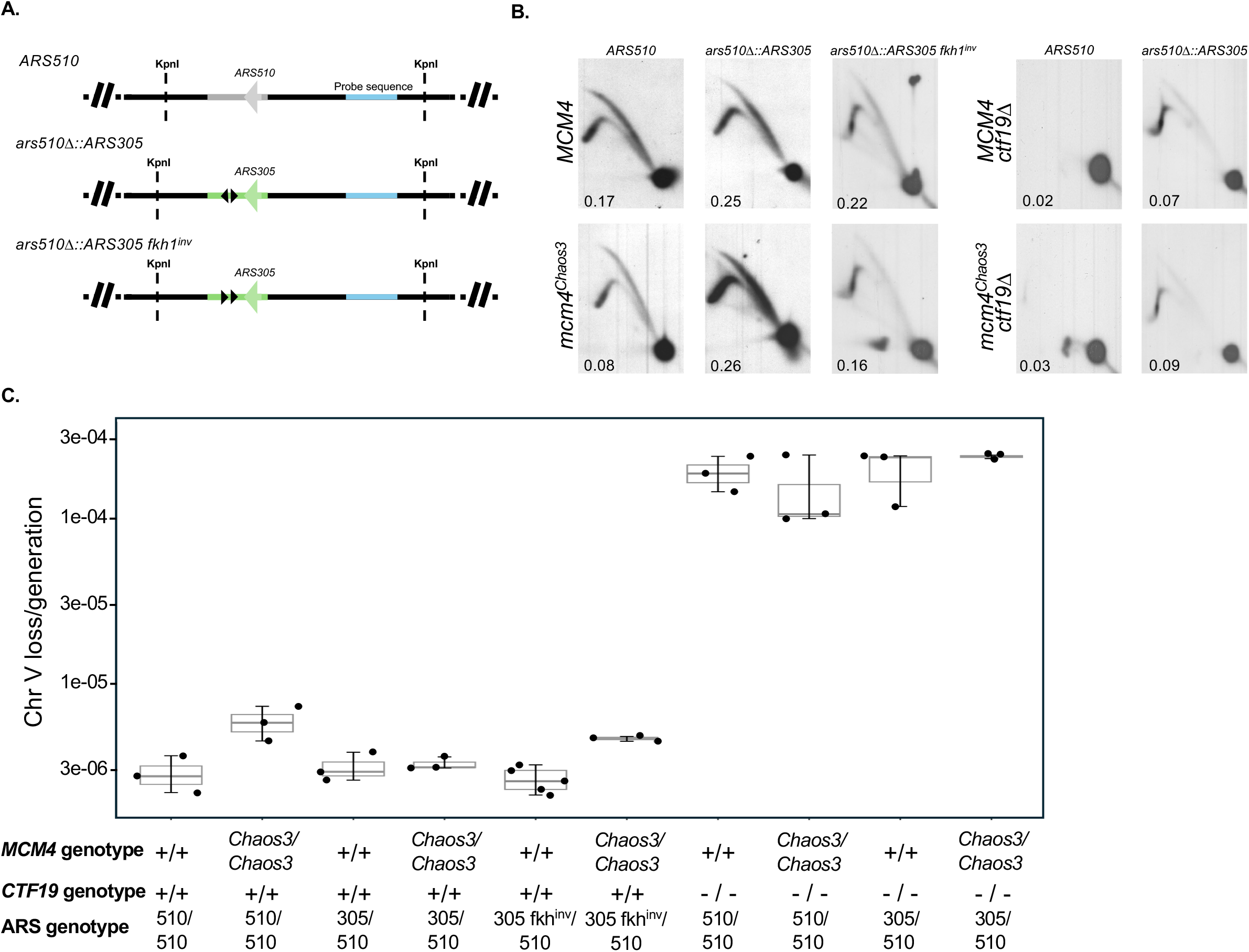
FKH 1/2 binding site requirement for origin replacement activity. (A) Diagrams of the region on Chr V containing the *mcm4^chaos3^*sensitive origin, *ARS510,* and the replacement of *ARS510* with *ARS305* creating the *ars510Δ::ARS305* mutant strain. The arrowheads indicate the orientation of the T-rich strand of the ACS. The restriction enzyme KpnI cuts this region of Chr V twice creating a ∼2.6 kb fragment used to assay origin activity at the ARS510 locus using 2D gel electrophoresis. The “probe sequence” (blue) was used to probe for the 2.6 kb fragment in the 2D gels (B). The 3’ FKH 1/2 binding site of the divergently oriented FKH 1/2 binding sites on *ars510Δ:ARS305* (black triangles) was inverted to create the *ars510Δ::ARS305 fkh1^inv^* strain. (B) Representative two-dimensional gel electrophoresis analyses of replication intermediates from the indicated strains in (A). Replication intermediates were analyzed in *MCM4* or *mcm4^Chaos3^* backgrounds, with or without *CTF19*, as indicated. The values represent the quantified signal associated with the replication structures in each sample. (C) Box and whisker plots showing the frequency of whole-chromosome V loss in homozygous diploid strains carrying either *MCM4* (+/+) or *mcm4^Chaos3^* (Chaos3/Chaos3) in combination with either *CTF19* (+/+) or *ctf19Δ* (-/-). The ARS genotype is indicated below each group for the origin that is present at the native *ARS510* locus. 510/510 represents a strain of yeast with two intact *ARS510* origins present in their native location while 305/510 represents a diploid with the dominantly marked copy of Chr V containing the *ars510Δ::ARS305* origin replacement and the other copy of Chr V containing native *ARS510*. 305 fkh^inv^/510 represents the corresponding *ars510Δ::ARS305 fkh^inv^* paired with *ARS510*.

We then performed 2D gel electrophoresis to determine origin activation of the various DDK recruitment mutants (Figure 7B). As shown previously, *ARS510* in a *mcm4^Chaos3^* background is reduced by half in comparison to the activity in wild type (Figure 7B, 0.17 vs 0.08). This defect in origin activity is rescued by the replacement of *ARS510* with *ARS305* (Figure 7B, 0.25 vs 0.26). In the *MCM4* background, inversion of the *ars510Δ::ARS305* FBS resulted in only a modest reduction in origin activity compared to the intact *ars510Δ:ARS305* (0.25 vs 0.22), consistent with the FBS contributing to but not being the sole DDK recruitment pathways active in a wild-type background. Critically, in the *mcm4^Chaos3^* background, inversion of the FBS substantially diminished the rescue of origin activity conferred by *ARS305* (0.26 vs 0.16), demonstrating that the FBS-mediated Fkh pathway of DDK recruitment is a major contributor to the rescue of origin firing under conditions of compromised MCM complex function.

To further test the contribution of each DDK recruitment pathway, we examined origin activity in a *ctf19*Δ background. Deletion of *CTF19* severely impaired *ARS510* origin firing in both the *MCM4* and *mcm4^Chaos3^* backgrounds (Figure 7B), consistent with ssDNA replication profiling and the dependence of this centromere-proximal origin on the Ctf19 pathway. Interestingly, the combination of *ctf19*Δ with the origin replacement strain, *ars510*Δ::*ARS305*, produced only marginal improvement in origin activity for both wild type and *mcm4^Chaos3^* (Figure 7B), indicating that functional FBS are not sufficient to completely rescue this initiation defect and that maximal origin activity requires both functioning DDK recruitment pathways.

We next assessed whether the observed changes in origin activity translate to changes in chromosome stability using our chromosome loss assay (Figure 7C). In a wild-type Ctf19 background, *mcm4^Chaos3^* elevated the chromosome loss rate approximately 3-fold compared to *MCM4*. Replacement of one *ARS510* allele with *ARS305* (305/510) suppressed this elevated loss rate back to near-baseline levels, consistent with the rescue of origin activity observed by 2D gels. However, when the FBS was inverted (305 fkh^inv^/510), chromosome loss was again elevated, demonstrating that FBS integrity is required for the rescue of chromosome instability in the *mcm4^Chaos3^*mutant and that the activity of this centromere-proximal locus is directly related to chromosome stability. In the *ctf19*Δ background, chromosome loss rates were dramatically elevated across all genotypes, approximate 40-fold above the wild-type baseline. Although *ars510*Δ::*ARS305* increased local origin activity slightly in the absence of Ctf19 (Figure 7B), this increase did not translate into any suppression of chromosome loss. Chromosome loss rates remained elevated in all *ctf19*Δ strains regardless of Mcm4 or ARS genotype, suggesting that the small amount FBS-mediated rescue of the *ARS510* locus firing is insufficient to overcome the chromosome instability caused by the loss of Ctf19. These data suggest that *ARS305* is almost entirely dependent on Ctf19 at the centromere regardless of the presence of its native DDK recruitment method, but Ctf19 alone cannot support *ARS305* origin activity in a *mcm4^Chaos3^* background, again exposing CEN V to replication delays and increased chromosome instability.

## Discussion

Our work establishes that the origin-specific defects associated with the *mcm4^Chaos3^* mutation extend beyond plasmid maintenance to chromosomal origin firing. By examining early S-phase origin activity we find that compromised MCM function preferentially reduces activity of a subset of centromere-proximal origins. We hypothesize the decrease in early S-phase origin firing at centromere-proximal origins results in a delay in centromere replication. This proposed defect in centromere replication is exemplified by an approximately three-fold increase in chromosome V loss—a chromosome harboring, *ARS510*, one of the *mcm4^Chaos3^*sensitive, centromere-proximal origins—a marginal increase in the test chromosome loss rate compared to the more extreme defect in origin firing we observed in the replication profiles. Centromeric DNA replication results in the dissociation of the kinetochore proteins from the centromere which is expected to disrupt kinetochore-microtubule interaction (25). Chromosome-microtubule interaction defects as a result of delayed centromere replication have been shown to activate the spindle assembly checkpoint, delaying chromosome segregation in response to unattached kinetochores (24, 36–38). It is likely that the activation of the spindle assembly checkpoint in response to our hypothesized delay in centromere replication in *mcm4^Chaos3^* mutants prevents higher rates of chromosome mis-segregation. In addition, replacement of *ARS510* with a robust, non-centromere-adjacent origin, *ARS305*, restores local origin activity and suppresses the elevated chromosome loss rates, providing a functional connection between robust centromere-proximal origin firing and faithful chromosome inheritance.

The origin replacement experiment demonstrates that proximity to the centromere alone does not account for sensitivity to the *mcm4^Chaos3^* mutation. *ARS305* retains robust activity at the *ARS510* locus; however, this robustness is not independent of chromosomal context. Inversion of a Forkhead Binding Site (FBS) diminishes the restoration of origin firing and eliminates the rescue of chromosome instability, whereas deletion of *CTF19* strongly compromises activity even when the FBS remains intact. Why is Ctf19 unable to fully support *ARS305* activity near *CEN5* in the *mcm4^Chaos3^* background? Hoggard et al. (39) described the outcome for origins tested for ORC binding *in vitro* and compared that to ORC binding *in vivo*. Three of sixty-six tested origins have low *in vivo* ORC binding but high *in vitro* binding, suggesting that chromatin can play a negative role in ORC binding *in vivo*. One of the three origins they identified is also an origin our study identified as being a *mcm4^Chaos3^* sensitive, centromere-proximal origin (Sup Fig 3, *ARS920*). If other centromere-proximal, *mcm4^Chaos3^*sensitive loci are located in similar “negative chromatin states” we speculate that in wild-type cells Ctf19 can overcome this negative chromatin state to activate the centromere-proximal origins, but Ctf19 is not able to fully overcome the competitive environment in a *mcm4^Chaos3^* background, resulting in the decrease in origin activity we observe. This negative chromatin context could also explain why *ARS305* is apparently unable to utilize its Fkh pathway at the *ARS510* locus in a *ctf19Δ* background. These results support a model in which kinetochore-dependent and Fkh-dependent DDK recruitment pathways cooperate to sustain robust firing at the replacement locus.

Shima et al. reported that the *mcm4^Chaos3^* mutation appears to destabilize the MCM4 protein (11). Protein blot analysis of mouse embryonic fibroblasts (MEF) revealed reduced amounts of MCM4 and MCM7 proteins in *mcm4^Chaos3/Chaos3^* MEFs (11). *Mcm4^Chaos3^* homozygosity also caused an approximately 60% reduction of all components of the MCM2-7 complex on chromatin compared to wild-type MEFs (40). In *Saccharomyces cerevisiae*, MCM proteins exist in vast excess with intracellular concentrations being 100-500 times the number of replication origins (41). This excess ensures proper initiation of all necessary replication origins (42). Like MEFs, reduced amounts of MCM proteins have been reported to compromise normal origin licensing and genome stability in yeast (42, 43). Reduction of MCM levels through the degradation of Mcm4 led to differential loading of the MCM helicase during G1, revealing differences in both origin sensitivity to a reduced MCM state and overall variability in MCM loading capacity at different origins (43). Because fluctuations in origin licensing capacity in a reduced MCM state would directly affect origin firing we compared our early S-phase origin activity profiles to Dukaj and Rhind’s MCM licensing data (43) and found a correlation between origins that have reduced firing in the *mcm4^Chaos3^* background and those that have reduced MCM loading as a result of Mcm4 degradation (Sup Fig 5, orange boxes). In addition, origins that retain wild-type origin firing levels or have an increase in firing in the *mcm4^Chaos3^* mutant maintain MCM loading in a reduced MCM state (Sup Fig 5, blue boxes).

One possible explanation for differential MCM loading in response to an overall reduction in MCM complexes is an origin’s affinity for the Origin Recognition Complex (ORC). In the nucleus, ORC recognizes and binds origins at the A and B1 elements (A/B1) (44–46), also referred to as the extended ARS Consensus Sequence (EACS) (47, 48). ORC subsequently recruits MCM-Cdt1 to origins and loads MCM onto the DNA. Cdt1 is then released, allowing ORC to load a second MCM complex by binding a less well defined and lower-affinity binding site called the B2 element (B2) (46). The presence of a B2, the orientation of the EACS compared to the B2, and the distance between the EACS and B2 are all determining factors of the affinity and efficiency of ORC MCM loading (49, 50). We did not find any obvious pattern relating EACS and B2 elements to the origins we identified as being sensitive, robust, or having improved firing in the *mcm4^Chaos3^*background.

Our early S-phase origin firing data demonstrated a variety of responses to a compromised MCM complex. As previously stated, we saw a significant subset of centromere-proximal origins that had reduced origin firing. Interestingly, we also observed origins that had an increase in firing in the presence of the *mcm4^Chaos3^* mutation. We hypothesize that the decrease in origin firing we observe at numerous loci throughout the genome increased the availability of limited replication factors that would normally in use by these origins. The increase in origin activity would then be a result of the redistribution of these limited factors. Why there is not a global increase in origin activity as a result of the newly available replication factors and what is directing replication factors to specific origins is unknown, but one could speculate that it could be due to origins having robust ORC MCM loading, the presence of Forkhead Binding Sites, chromatin accessibility, or any combination of the proposed mechanisms.

To assess origin-specific sensitivity to the *mcm4^Chaos3^* mutation without the potential compensatory effects of neighboring origins, we measured the loss of single-origin plasmids containing either *ARS510* or *ars510Δ::ARS305*. The combination of plasmid loss rates determined by Southern blot and by plating allowed us to distinguish the contributions of replication and segregation defects to plasmid instability. While *mcm4^Chaos3^* increased replication-based loss of both plasmids, the plasmid containing *ars510Δ::ARS305* remained more stable than its *ARS510* counterpart and supports the finding that origins differ in their *mcm4^Chaos3^* sensitivity. Notably, segregation defects contributed substantially to the loss of the *ARS510* containing plasmid in *mcm4^Chaos3^* cells, whereas replication defects accounted for most of the loss of the *ars510Δ::ARS305* containing plasmid. These findings suggest that features associated with *ARS305* promote robust plasmid maintenance in the presence of a destabilized MCM helicase.

Overall, this work illustrates how defects in MCM function may specifically disrupt the robust and timely replication of centromere DNA, and how such replication defects may in turn compromise kinetochore integrity, faithful chromosome segregation, and the preservation of genome stability.

## Materials and Methods

### Yeast Strains and Culture Conditions

Strain HM14-3a (A364a background; *MATa, CDC7, bar1, trp1-289, leu2-3,112 his6, ura3-52, can1-100* (51); and its derivatives were used to construct all strains for this study. A derivative of HM14-3a with the functional copies of *URA3* and *CAN1* genes was used to construct additional “marked chromosome V” strains for fluctuation analysis. A complete strain list can be found in (Strain list). We verified each transformation by PCR, Sanger sequencing, and resolving whole chromosomes on a CHEF gel to check for normal rDNA array size on Chromosome XII. Primers used to amplify markers with homology can be found in (Primer list).

To minimize false positives during fluctuation assays due to *de novo* point mutations in the large *CAN1* gene, we inserted an additional copy of *CAN1* on the right arm of Chr V to create the “marked chromosome V” strains used for fluctuation analyses. First, antibiotic selection marker NAT was amplified from pFvL99 with homology arms to replace a 99 bp fragment downstream of *SIT1* on the Watson strand near the native copy of *CAN1* (primers 1 & 2). An additional copy of *CAN1* was then introduced into the *SIT1::NAT1* strain in the opposite orientation on the opposite arm of Chromosome V near *UBP9* (primers 3 & 4).

Standard lithium acetate transformation with a fragment amplified from pRS400 (*KanMX*; Kanamycin/G418 resistance marker) with homology to the flanks of *CTF19* was used to make the *ctf19Δ* gene replacement (*ctf19Δ::KanMX*) (primers 7 & 8). Transformants were selected on plates containing G418.

Strains with *ARS305* in place of *ARS510* (*ars510Δ::ARS305*) were created using the CRISPR plasmid pYCpGal (52) with guide RNA targeting *ARS510* (primers 11 & 12) and an *ARS305* template amplified from genomic DNA with homology targeting the region immediately surrounding *ARS510* (primers 9 & 10). A 653 bp fragment containing *ARS305* replaced 421 bp containing *ARS510*, with *ARS*s oriented in the same direction in each fragment and positioned approximately within the center of the fragment.

The 3’ Fhk1/2 consensus binding sequence of *ARS305* was reversed and one base pair was deleted to create *ars510Δ::ARS305^fkh^ ^inv^*. These strains were created using the CRISPR pYCpGal plasmid with guide RNA targeting *ARS305* (primers 16 & 17) and a Fkh1/2 inversion template containing the 7 bp mutation and homology to the surrounding region (primers 13-15). Native *ARS305* and *ARS305* at the replacement location on Chromosome V were screened for the presence of the Fkh inversion. Strains were only kept if the inversion was present at the replacement location and the native *ARS305* Fkh1/2 consensus binding sequences remained intact.

Plasmids used in the plasmid loss assay were constructed by PCR amplifying a ∼1.2 kb fragment from chromosome V containing either *ARS510*, *ARS305*, or *ARS305* fkh^inv^. The DNA fragment was inserted into a *CEN5*-containing derivative of pRS406 which contains ampicillin resistance gene Amp^R^, ColE1/pMB1-derived origin *E. coli* origin, *CEN5*, and *URA3*.

Yeast strains were grown in complete synthetic media with 2% glucose unless otherwise noted. When possible, for experiments comparing different strains, samples were collected in the same media.

### Southern blotting

DNA from each gel was transferred to a GeneScreen Hybridization membrane (Revvity Health Sciences) using standard Southern blotting protocols. We then hybridized each sequence of interest using a ^32^P-labeled probe. The blots were exposed to X-ray film and to Bio-Rad Molecular Imaging FX phosphor screens for visualization and quantification of signal intensity.

### Flow cytometry

Cells for flow cytometry were fixed in 70% ethanol before processing for flow cytometry. Fixed cells were washed with 50 mM sodium citrate, sonicated, and resuspended in 500 μL 50 mM sodium citrate. RNase A was added to a concentration of 0.25 mg/mL and the samples were incubated for 1 h at 50°C. Proteinase K (50 μL of 20 mg/mL) was then added, and cells were incubated another hour at 50°C before staining with 1 μM Sytox Green. Cells were analyzed on a BD Canto II flow cytometer and flow cytometry data was analyzed using FlowJo software.

### Cell collections in hydroxyurea

Cells were grown in logarithmic phase to a culture density of OD_660_ = 0.25. Cultures were arrested in α-factor for 1.25 doublings, then the culture was split to remove approximately 25 mL of α-factor arrested cells for flow cytometry/budding analysis to ensure that cells would undergo a normal S phase in the absence of Hydroxyurea (HU). HU was added to the remaining culture to a final concentration of 200 mM; 30 minutes later both cultures were released into S phase by addition of Pronase (0.15 mg/mL final concentration). 100-300 mL was collected for the following times post-release: 15, 30, 60, 90, and 120 minutes. For single stranded assays, 250 mL α-factor arrested samples were collected immediately before release to assess single strandedness during G1 phase. Collection vessels contained frozen pellets of 0.1% sodium azide in 0.2 M EDTA to halt growth. A small sample was taken for flow cytometry after collection, and the remaining dry cells were pelleted by centrifugation, washed in cold glass-distilled water and stored at −20°C until preparation for 2D gel electrophoresis and single strand assays.

### 2-Dimensional agarose gel electrophoresis

For each strain of interest, a time course of cells was collected as described above: 300 ml culture volume was collected for each time point and chilled on frozen sodium azide and EDTA. Genomic DNA was purified using a “Zymo-NIB (nuclear isolation buffer) & Grab” DNA purification protocol (53). The DNA purified from the 300 ml of culture was digested with the enzyme KpnI for 5 hrs at 37° C to analyze *ARS510* (2.6 kb fragment). The first-dimension agarose gel (0.5%) was run at 1 V/cm for 18 hours without ethidium bromide. The 1.1% agarose second-dimension gel with 0.3 µg/ml ethidium bromide was run in the cold room with buffer circularization at 7 V/cm for 4-5 hour then Southern blotted and probed for the region of interest using a ^32^P labeled *ARS510* probe (primers 18 & 19). Because cells were collected in HU, replication activity was measured as the hybridization intensity (determined using a BioRad Personal Molecular Imager) from both the bubble and Y arcs as compared to the 1N spot, as replication in Hydroxyurea is slowed such that all signal results from active replication at the region of interest (54).

### Single stranded DNA replication assay

To extract DNA for single stranded assays, cells collected as above were embedded in three 90 μL 0.5% SeaPlaque agarose plugs and prepared as CHEF gel plugs. Plugs from S phase time samples were labeled *in gelo* with with Cy3-dUTP, while plugs from the corresponding G1 samples were labeled with Cy5-dUTP. Labeled DNA was purified and hybridized to Agilent ChIP-on-chip 4×44 *S. cerevisiae* microarrays as described (55). We processed the array d ata essentially as described (53): we eliminated outliers of high S/G1 values by filtering out values more than 8 standard deviations above the mean and likewise eliminated low outliers (values less than half the baseline, set as the mode of the distribution of S/G1 values). We then applied LOESS smoothing over a 10 kb window to get smoothed data points at 0.25 kb intervals and re-scaled the data between 1 and 2 as described (53). Finally, to avoid false values due to sub-telomeric repetitive sequences, we trimmed off the terminal 10 kb of each chromosome. We used data from the 60’ S phase samples in this work. Two biological replicates of this experiment were performed. To facilitate comparison of wild type and *mcm4^Chaos3^*ssDNA replication profiles, the absolute difference between the means of the smoothed and scaled wild type and *mcm4^Chaos3^* S/G1 values were converted to Z-scores (units of standard deviation) as described (51). After multiple testing correction (56) on the basis of a false discovery rate of 0.05 the threshold for significance was set at Z = 3.447. The processed ssDNA data and Z-scores are available in (Figures data). Raw data will be deposited in GEO upon publication.

### Replication profiling by sequencing

G1 and S phase (30, 60, 90 min) samples were collected from cells released from G1 into S phase as above. DNA was extracted from frozen cell pellets using modified version of the Hoffman-Winston protocol (57). Cell pellets were resuspended in 200 µL lysis buffer (2% Triton X-100, 1%SDS, 100 mM NaCl, 10 mM Tris (pH 8), 1 mM EDTA), and 200 µL phenol/chloroform/isoamyl alcohol (25:24:1). 300 mg of acid-washed glass beads (∼500 micon size) were added to the resuspended cells and the mixture was vortexed for 8 minutes. After vortexing, 200 µL of Tris-EDTA (TE) buffer was added and the mixture was centrifuged at top speed. The aqueous layer was removed and added to 1mL room temperature 100% ethanol and centrifuged for 2 minutes top speed. The supernatant was decanted and 400 µL of TE buffer + 30mg RNaseA was added to remove RNA contamination. The samples were incubated for 30 minutes at 37°C. After incubation, 10 µL of 4M ammonium acetate and 1 mL of room temperature 100% ethanol was added to the sample and mixed by inversion. The sample was once again centrifuged at top speed for 2 minutes. The supernatant was decanted and the residual ethanol air dried. The final pellet of DNA was resuspended in 50 µL of water. The DNA was then cleaned and concentrated using the DNA Clean & Concentrator kit from Zymo Research (Cat. No. D4004) before the preparation of sequencing libraries. Libraries were prepared using the Illumina DNA Prep kit (Cat. No. 20025520) and sequenced as paired-end reads (58) on a NextSeq 2000. Reads were aligned to the *S. cerevisiae* S288C reference genome (sacCer3/R64-1-1) with BWA-MEM and duplicate reads were removed using samtools. Per-base read depth was then obtained from reads with a mapQ score of at least 60. Mean read depths were calculated for bins of 1 kb with a step size of 100 bp. For each S phase sample, a raw S/G1 ratio was computed for each bin. These values were then normalized as follows. Because the S phase was proceeding in HU, the bulk of the bins will not yet be replicated and therefore the “true” baseline S/G1 ratio should be 1. To normalize the raw ratios to this baseline, the mode of the distribution of ratios was obtained as described for the ssDNA assay above. The raw ratios were divided by this value to set the baseline for the replication profiles. LOESS smoothing was applied with a window size of 16 kb and finally the terminal 10 kb of each chromosome were trimmed to eliminate subtelomeric repeated sequences. Processed S/G1 ratios are available in (Figures data). Raw reads will be deposited in NCBI SRA upon publication.

### Fluctuation assay

Fluctuation assays were performed by the method described by G. Lang (59), with modifications described in Jiang et al. (60). Additional modifications to the protocol described to adjust for mutation rates in the strains used in this study include: the addition of ClonNAT to selective media on Day 1 to select for the engineered marked chromosome V in this study, an initial dilution of 1:5000 of saturated culture to inoculate wells, final plating on 15cm (large) *SC-Arg-Ser+Can+5FOA* plates, bringing well volume to 100ul using sterile media before plating to minimize background. Loss rates were calculated using the R package rSalvador (61).

### Plasmid loss assay

Quantification of plasmid loss for different strains was performed in two parts: Southern blotting to assess plasmid loss due to replication errors and standard selective and nonselective plating to assess total plasmid loss. Strains of interested were transformed with 1 of 2 different plasmids: p510-CEN or p510.305-CEN. A single colony from each plasmid harboring strain was picked and grown in selective conditions in SC-ura3 liquid medium for 10 doublings at 30 °C with rotation (∼48 hours). The saturated, selective culture was then diluted 1:1000 in SC + 2% glucose and again grown to 10 doublings in the same conditions as above. DNA was extracted from both the selective and nonselective growth cultures via a “Smash-&-Grab” protocol (57). The dried DNA pellets were dissolved in TE/RNase A (5µg/ml) overnight before overnight digestion with Pst1-HF and run on a 0.7% agarose gel in 1x TBE, 0.3 µg/ml EtBr for ∼6 hours at 95v. The same cultures that were grown for DNA extraction were used fo r the plating portion of the plasmid loss assay. Serial dilution of each culture was performed in -N and 100 µL were plated on SC and SC-ura3 agar plates yielding 30-300 countable colonies per plate. Plasmid loss rates were then calculated based on the number of colonies on selective vs. non-selective plates and the known number of generations of growth in non-selective medium.

### Co-culture experiment

Wild type and *mcm4^Chaos3^* that were otherwise isogenic were grown to saturation in synthetic medium with 2% glucose and mixed in equal proportions in two biological replicate pairs. The mixed culture was diluted 1:1000 in synthetic medium with 2% glucose (for a culture volume of 5 mL) and grown at 30°C for 9-10 population doublings. The cycle of dilution and growth was repeated twice more, for a total of ∼28 population doublings. DNA was extracted from samples taken from the initial and the final culture and the allele distribution of *MCM4* vs. *mcm4^Chaos3^* in each sample was determined by sequencing (Plasmidsaurus) of PCR amplicons (primers 20 & 21), with a minimum of 4,500 reads per sample.

## Acknowledgements

We are grateful to Steven Lee, Paula Levan, and Batool Mutawe for technical assistance and strain construction. MJD holds the William H. Gates III Endowed Chair in Biomedical Science.

## Supplemental Figure Legends

**Supplemental Figure 1:**
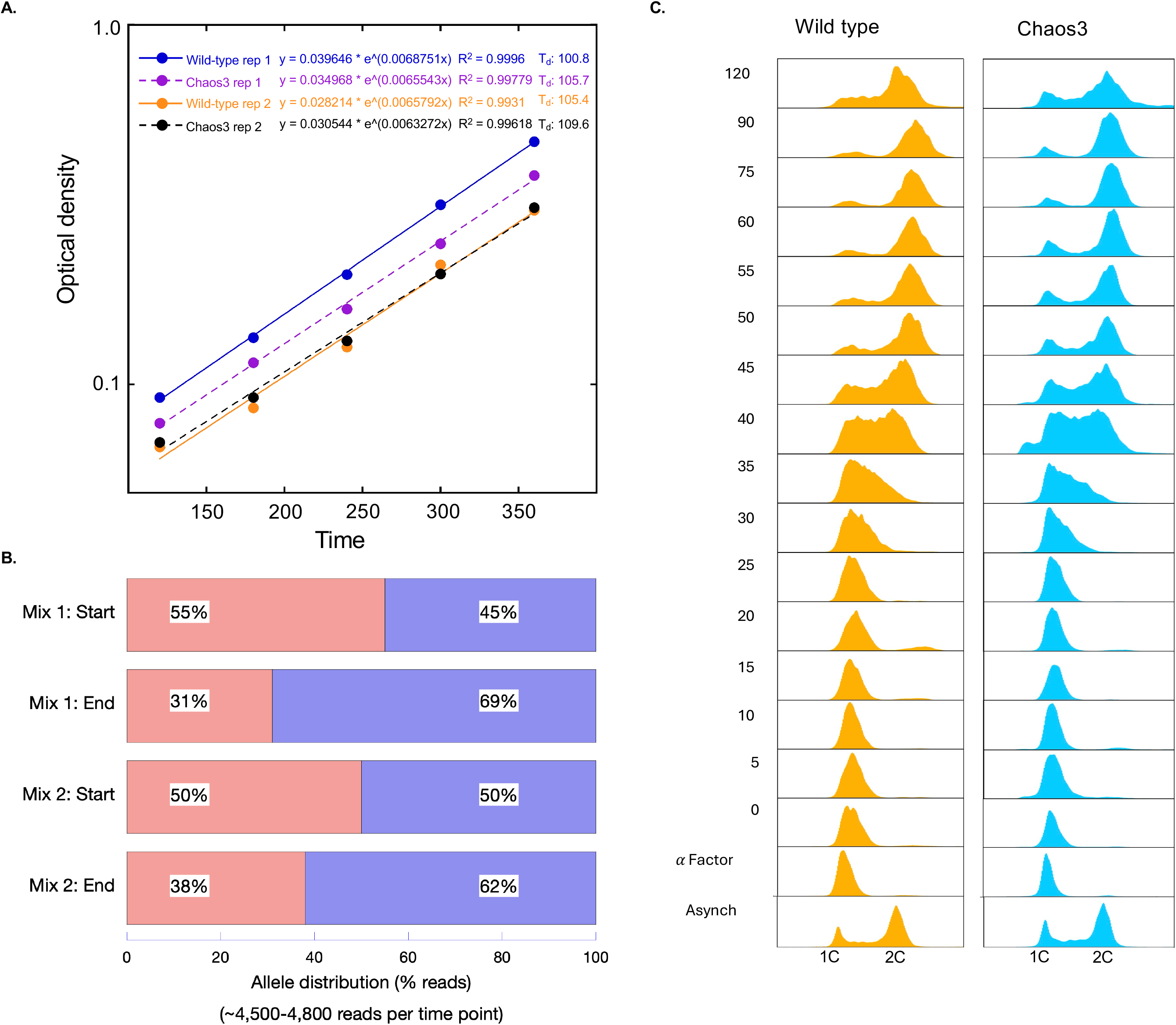
Growth rate and S-phase progression comparison of *MCM4* and *mcm4^Chaos3^* cells. (A) Growth curves of two wild type (blue and yellow points) and *mcm4^Chaos3^* (purple and black points) biological replicates. Optical density measured over time plotted on a logarithmic y-axis. Solid (wild type) or dotted (Chaos3) line for each replicate in their respective color represents line of best fit with exponential growth equation and R^2^ value for each replicate. Estimated doubling times are 100.8 and 105.4 minutes for wild type and 105.7 and 109.6 minutes for Chaos3. (B) Flow cytometry time course tracking genome duplication of *MCM4* (Wild type, orange) or *mcm4^Chaos3^* (Chaos3, blue) cells. Delay in S-phase progression of *mcm4^Chaos3^* cells can be seen starting at 40’ as indicated by the volume of cells in 1C versus 2C when compared to the same wild type sample. (C) Competitive growth assay confirms a slight growth advantage (2.5 minutes) of wild type over *mcm4^Chaos3^*.

**Supplemental Figure 2:**
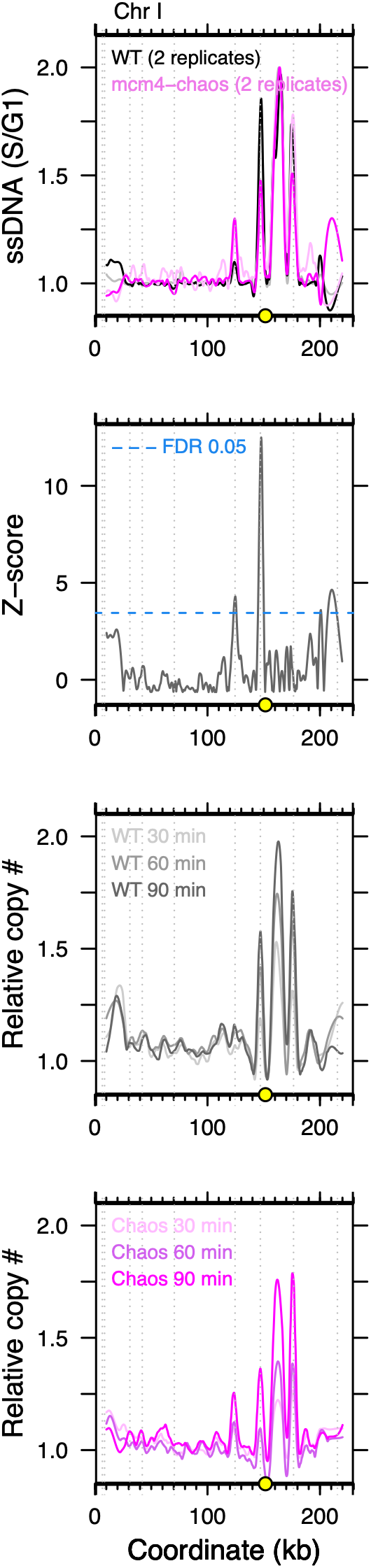
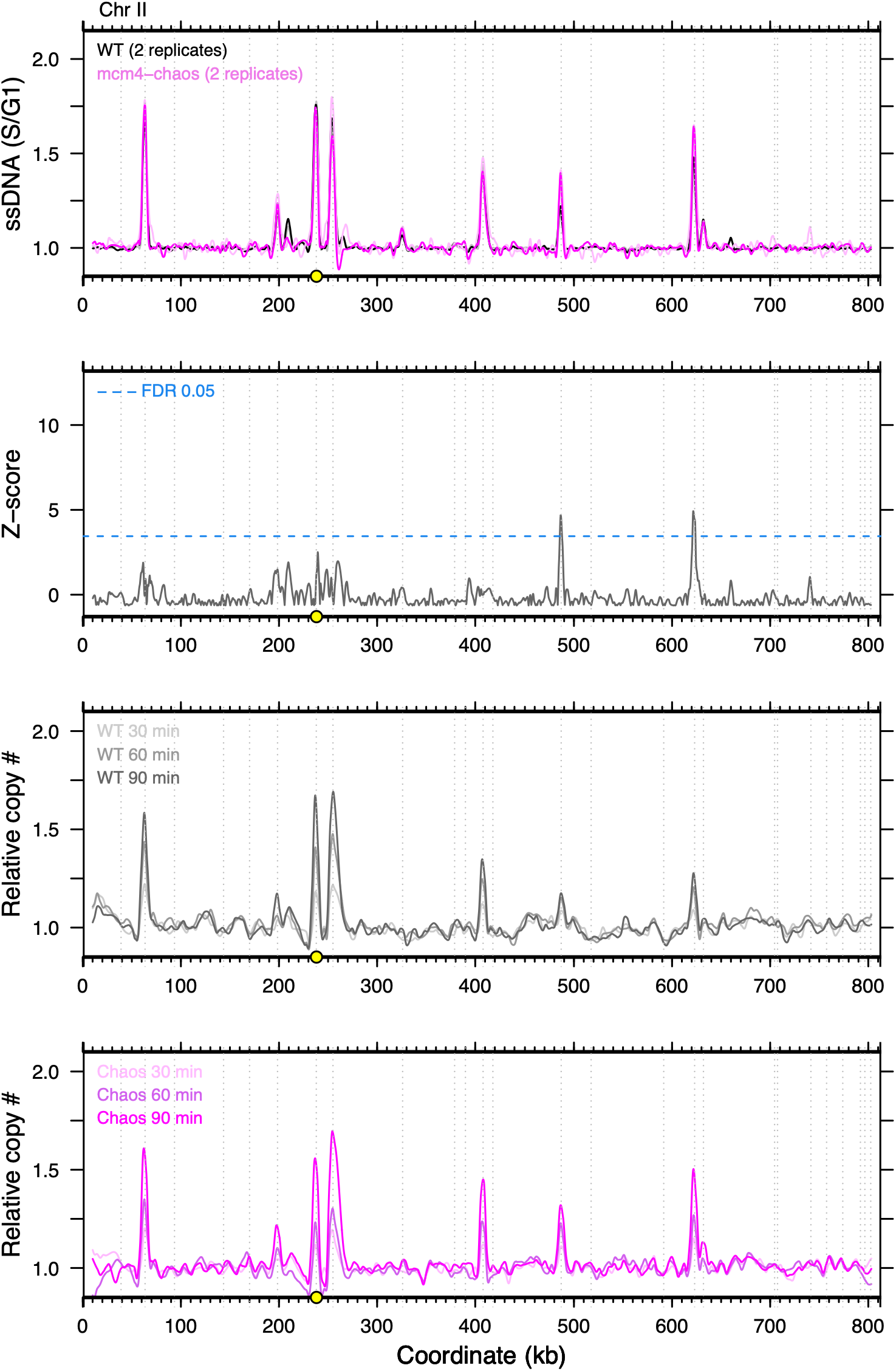
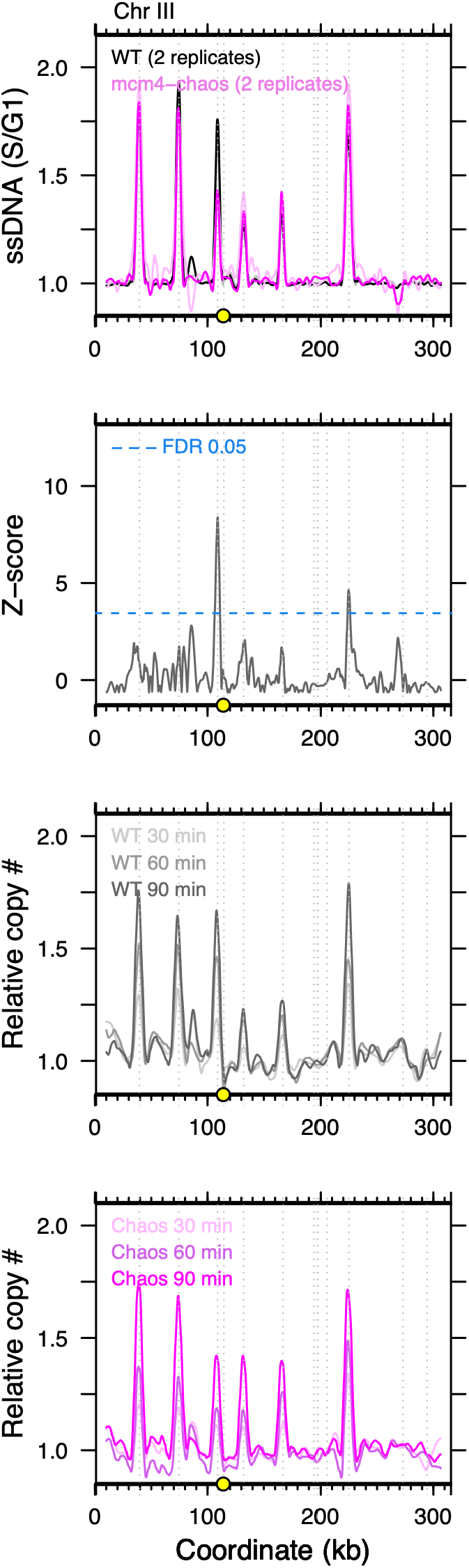
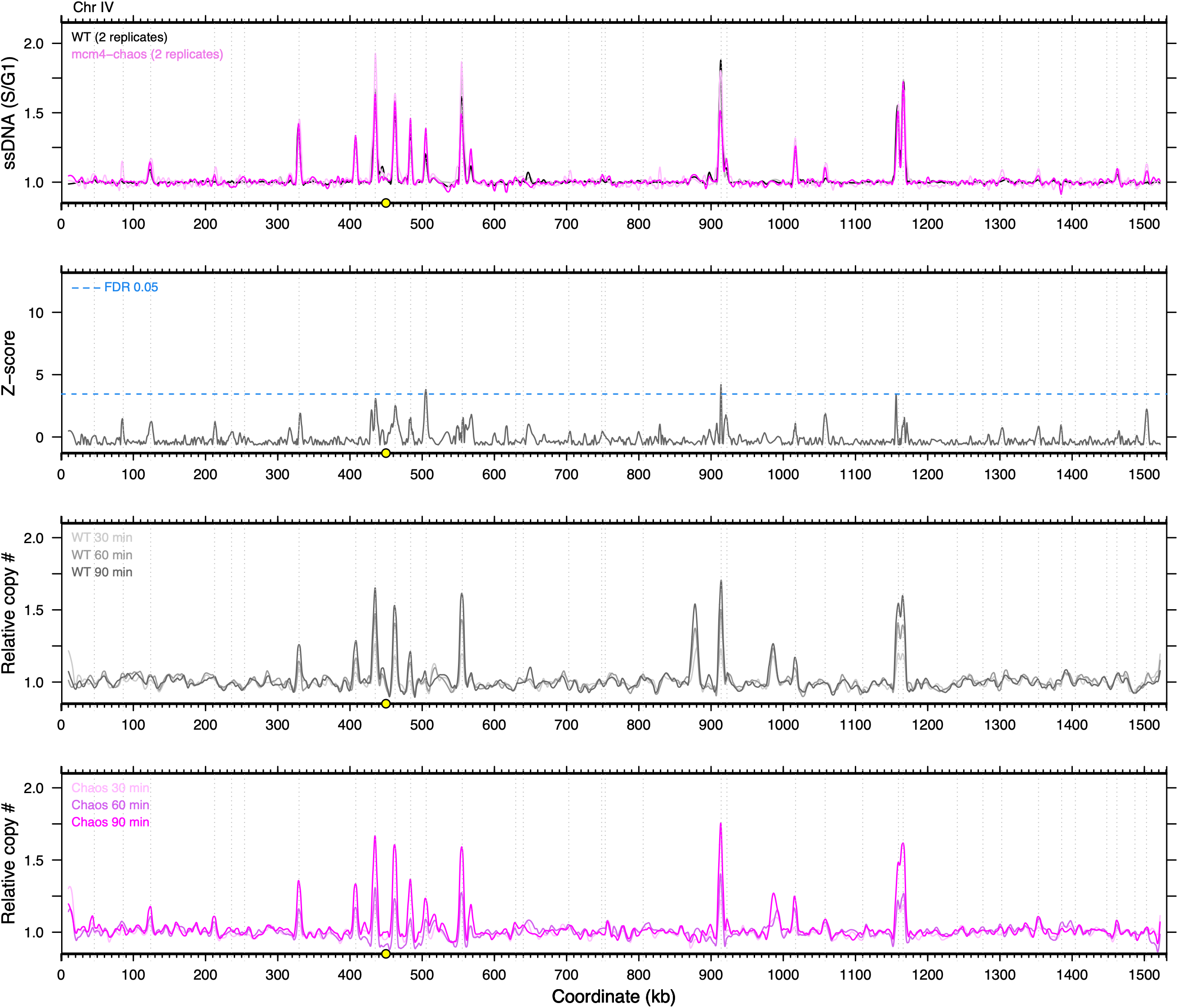
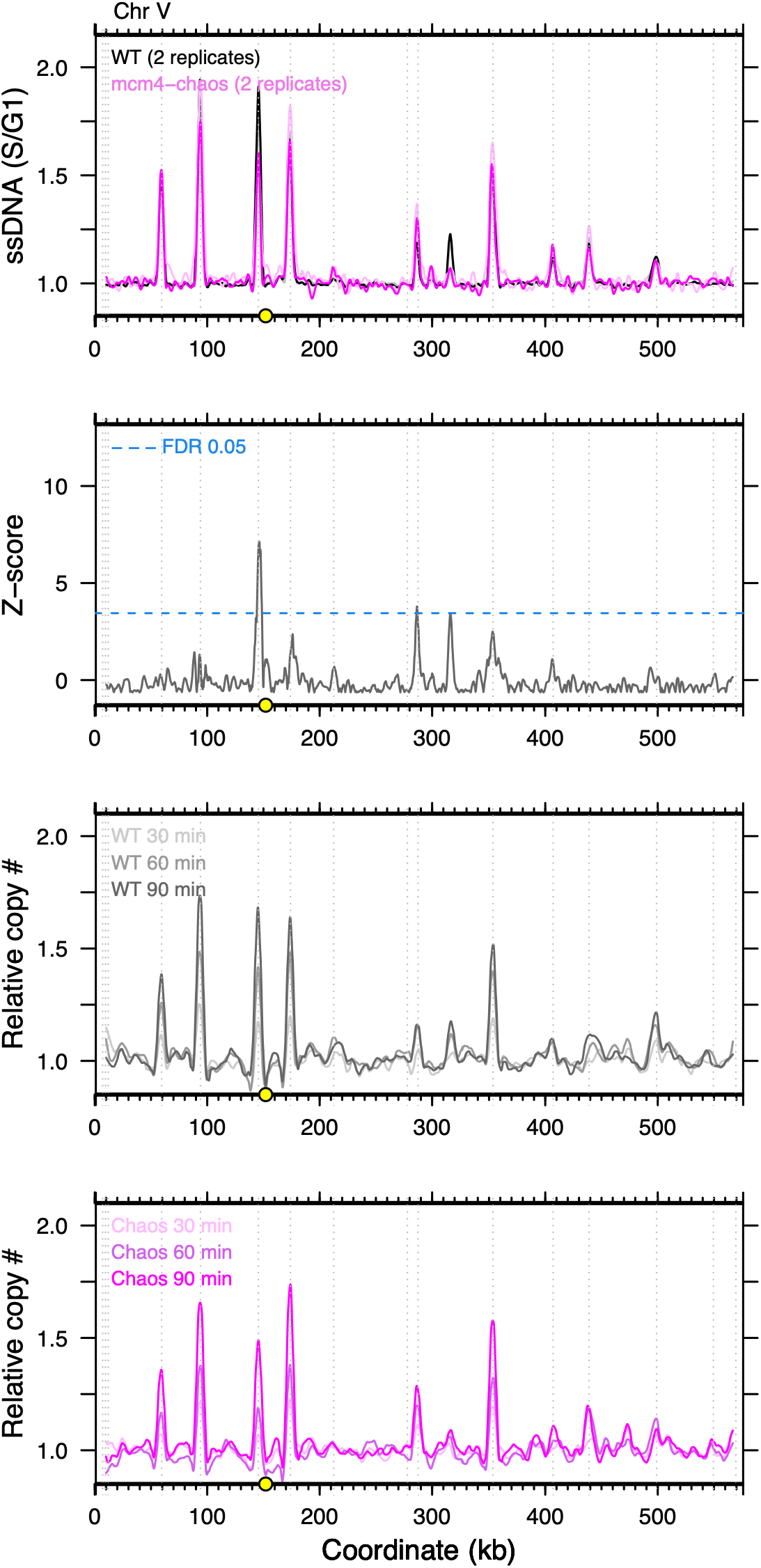
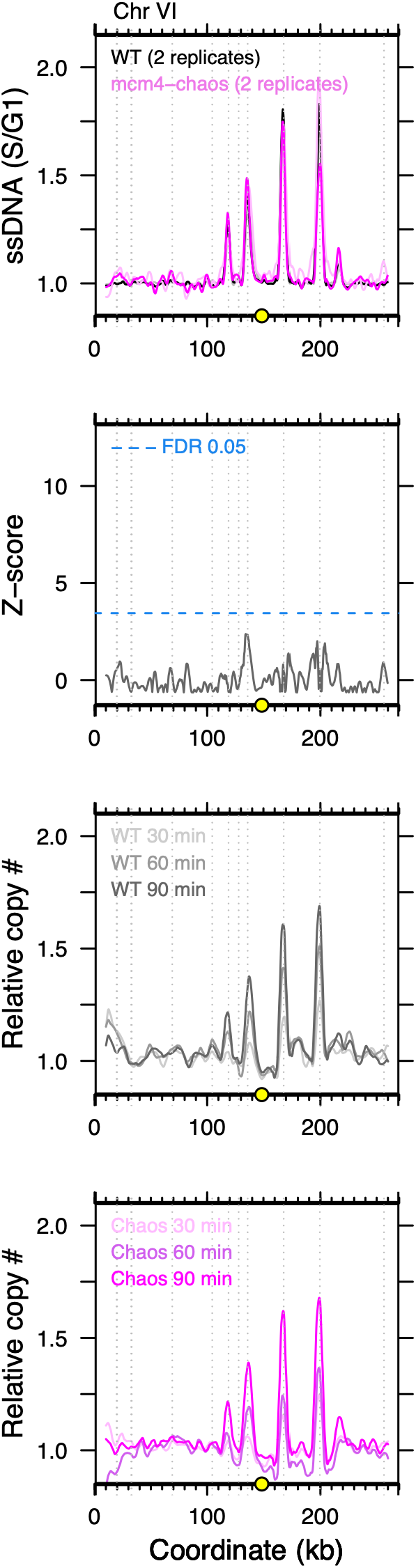
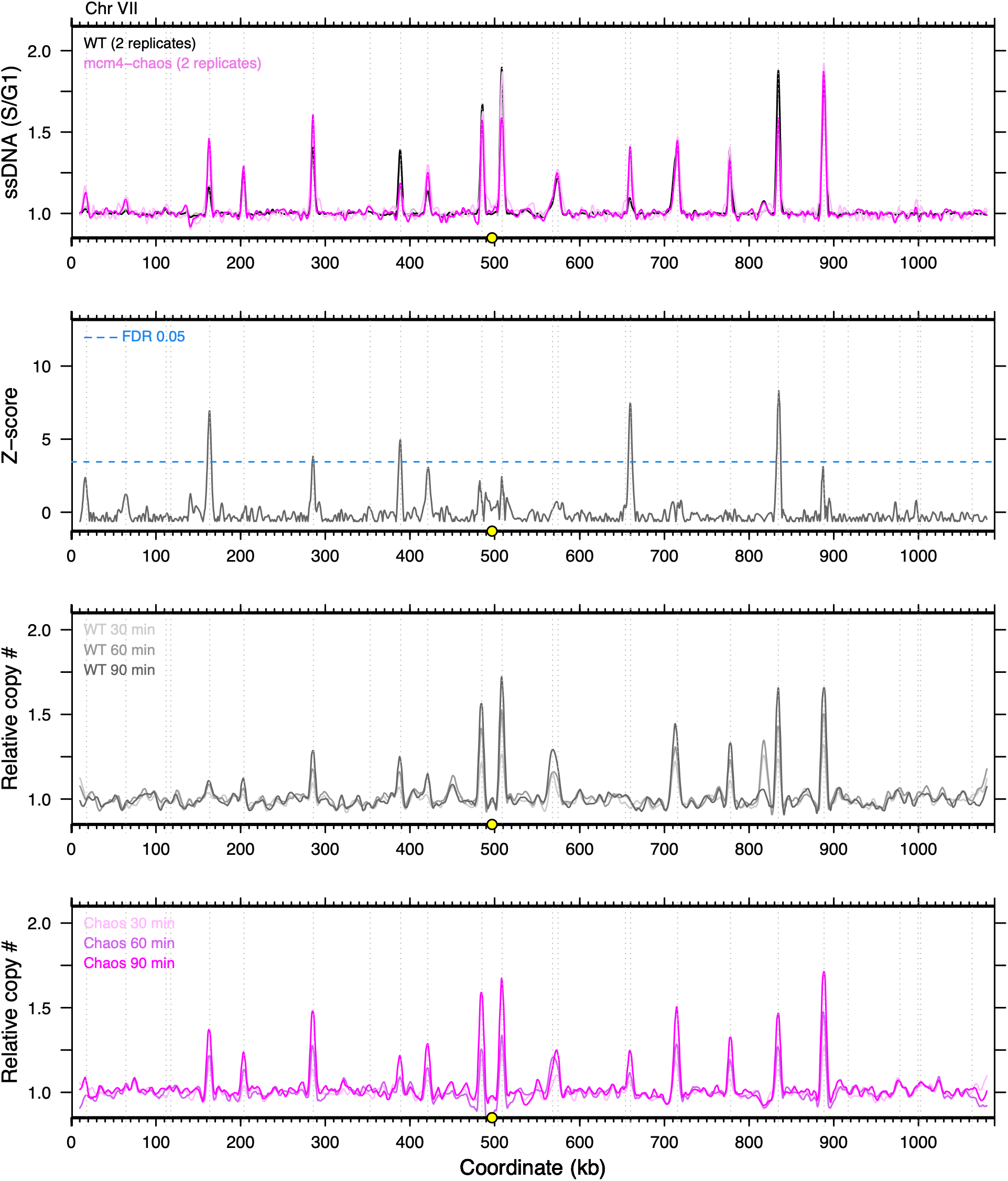
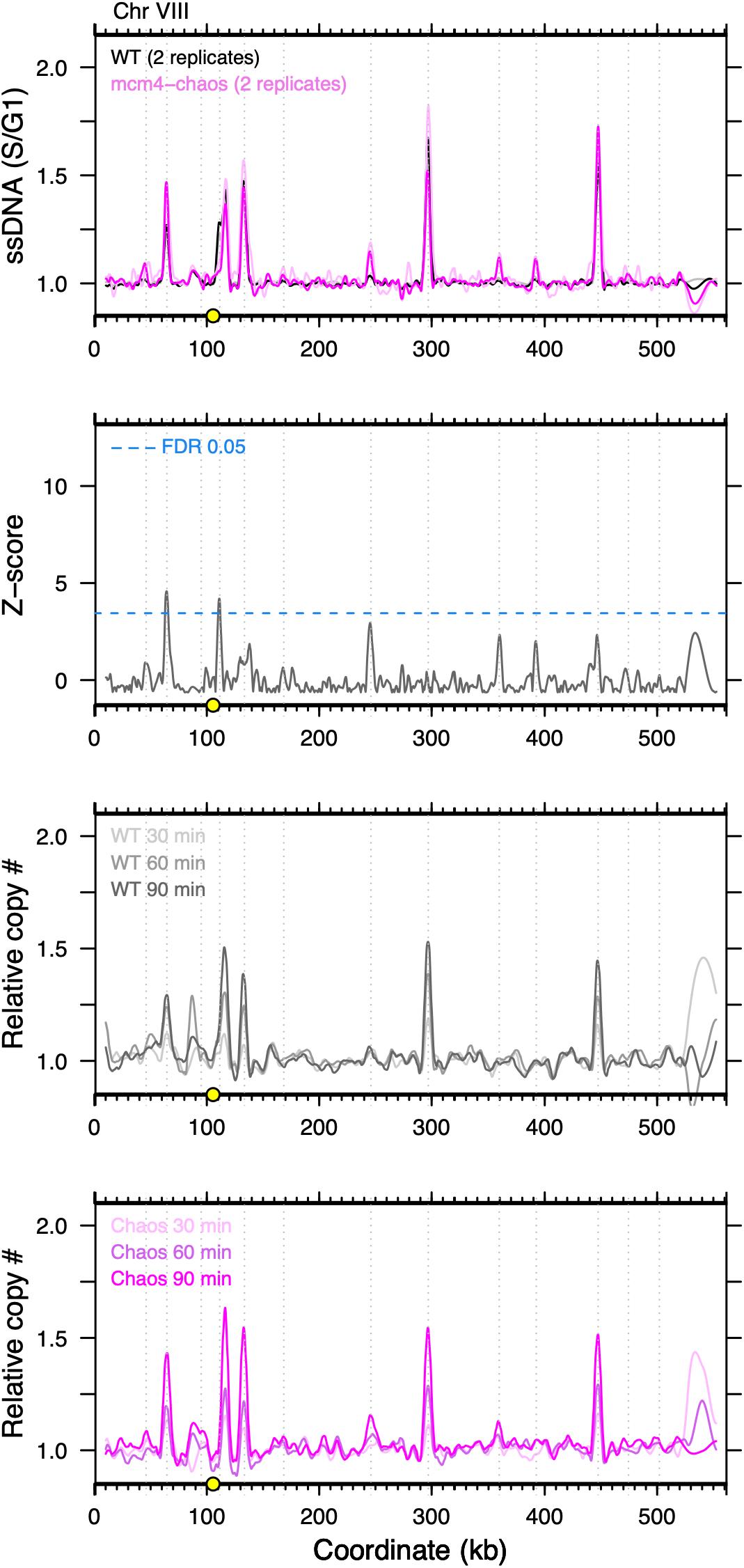
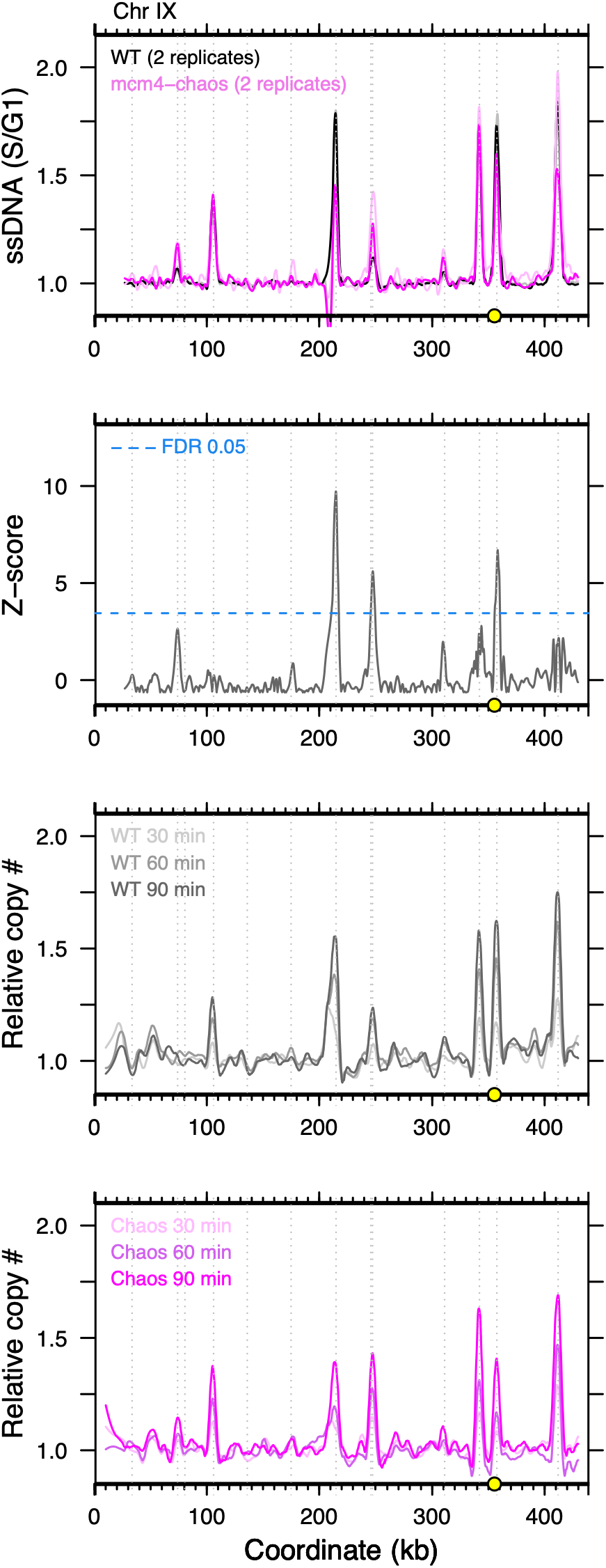
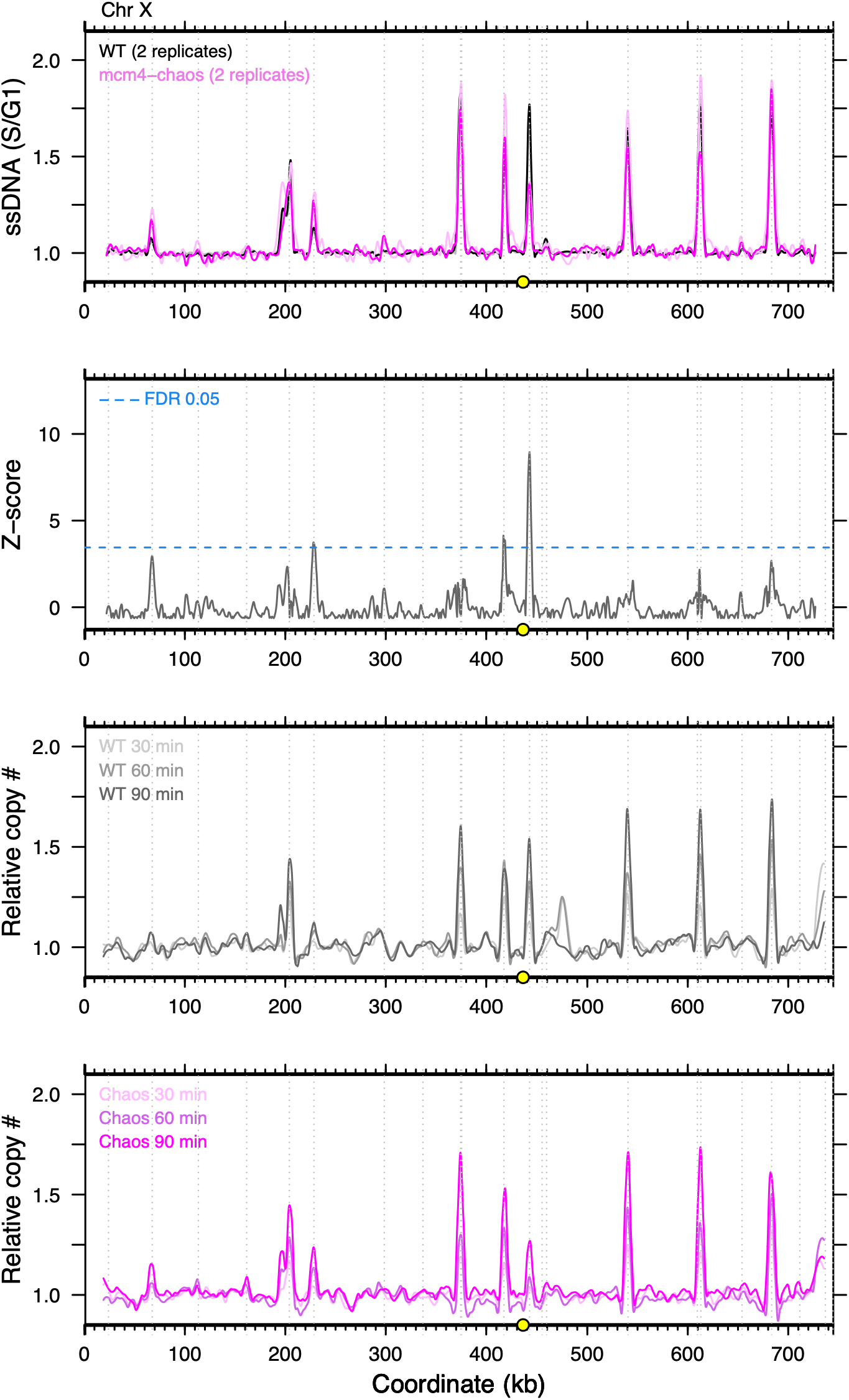
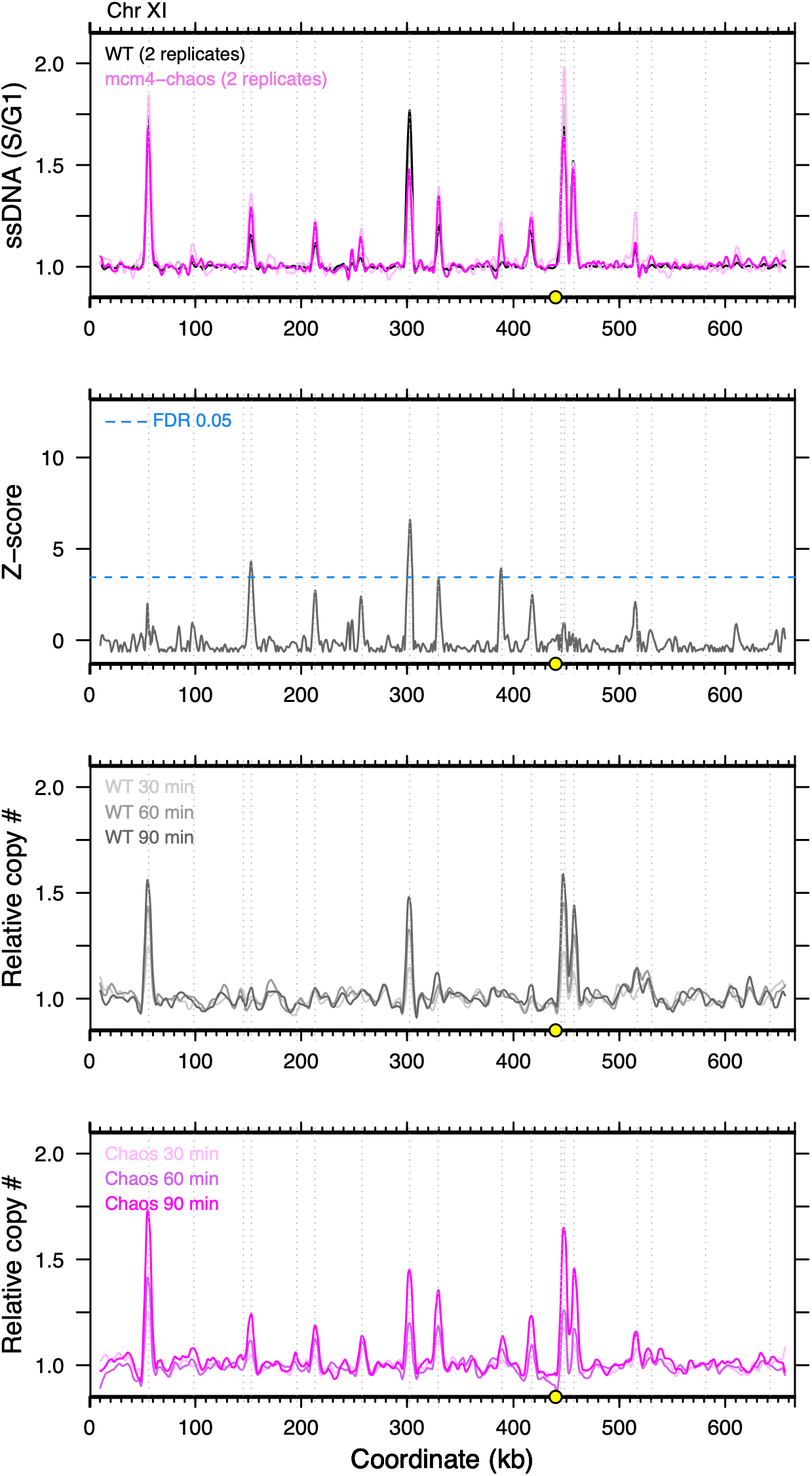
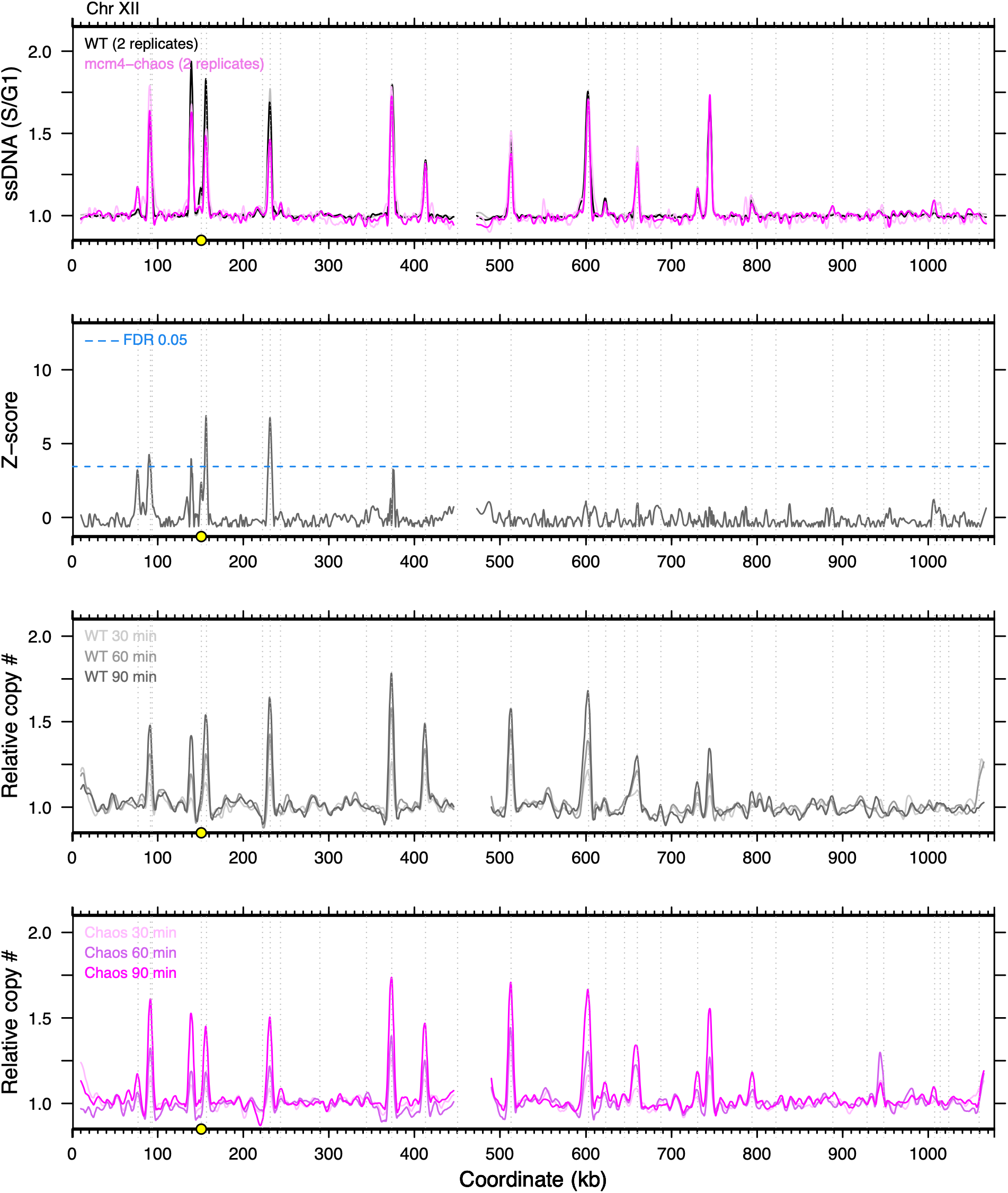
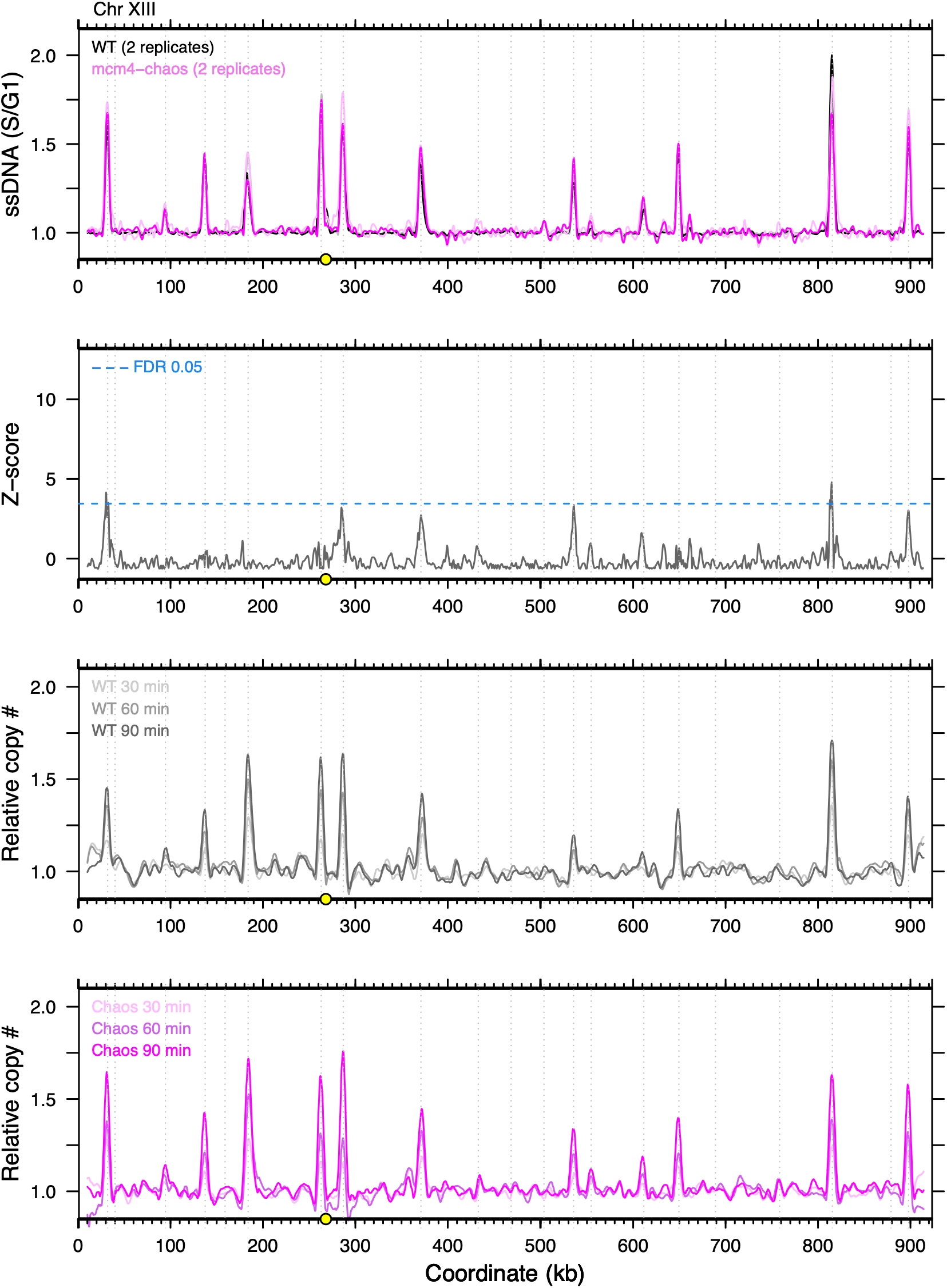
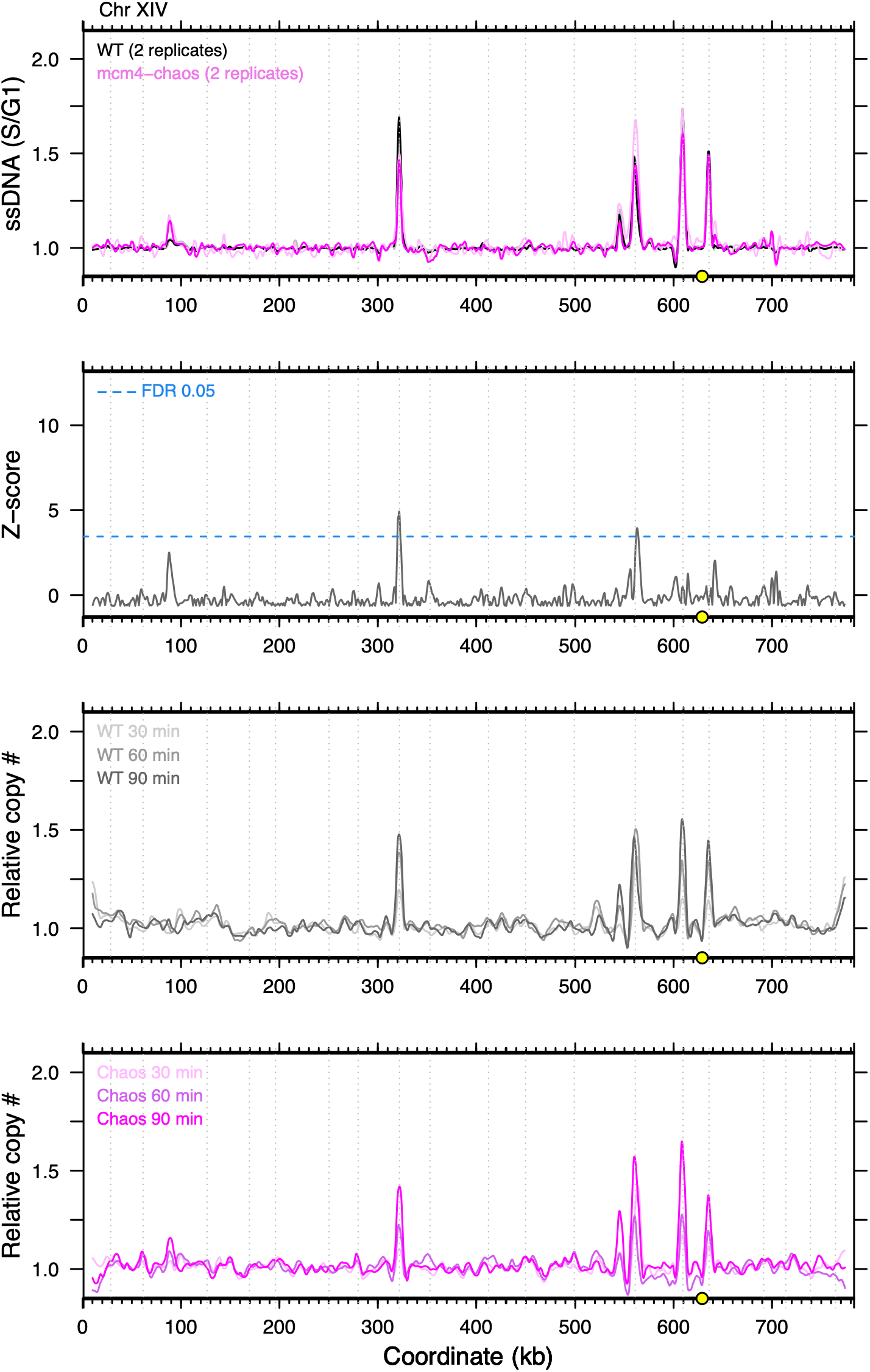
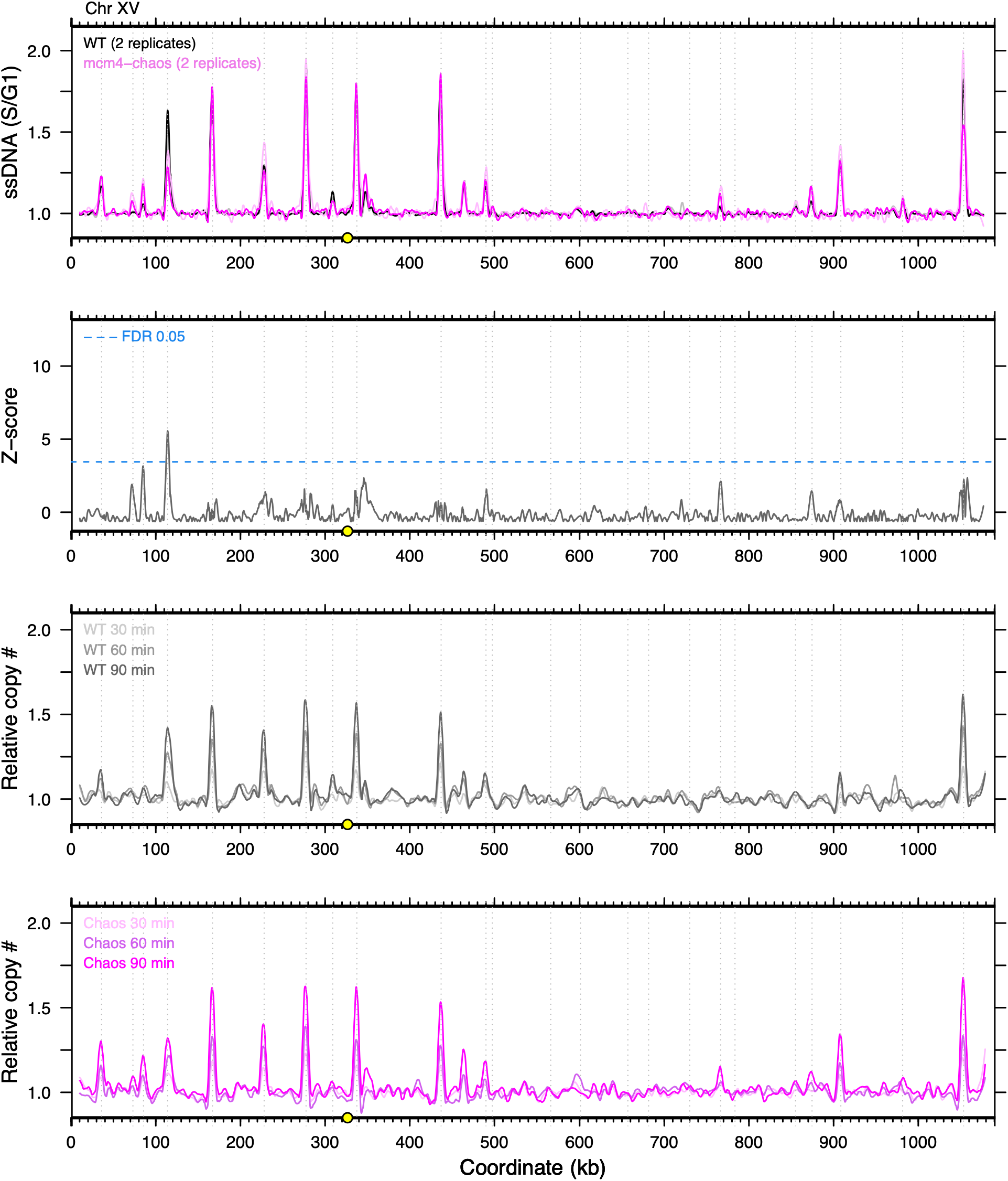
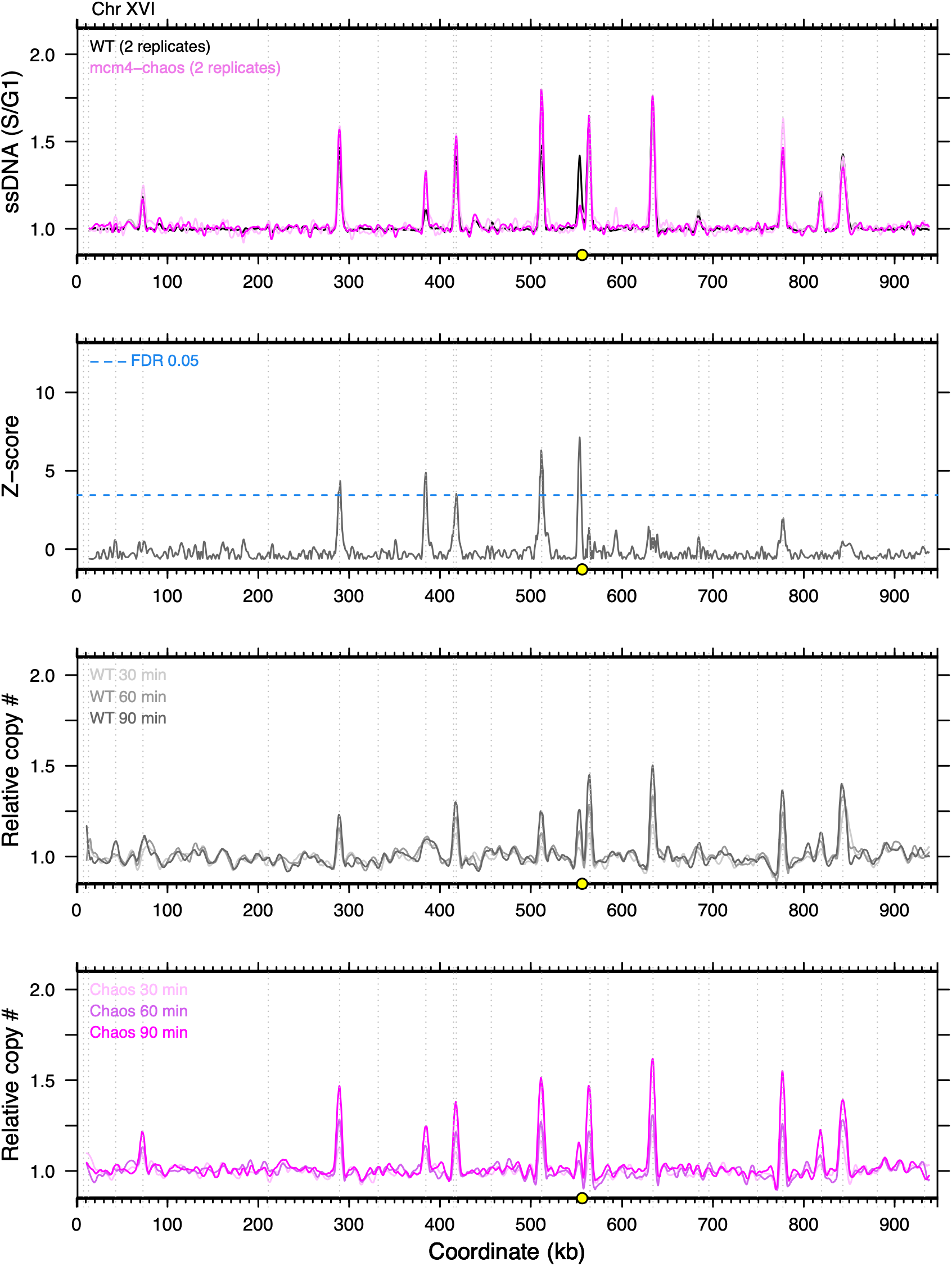
Replication profiles for each chromosome measuring early S-phase origin activity. (A) Each chromosome consists of four different panels. The x-axis for every panel corresponds to chromosomal position (kb) and the centromere is indicated by a yellow circle. Vertical dashed lines indicate locations of confirmed origins from OriDB (62). From top—Panel 1: Measurement of ssDNA in S phase relative to ssDNA in G1. The data were then overlaid for comparison of origin firing of two wild type biological replicates (black and gray lines) and two *mcm4^Chaos3^* replicates (light and dark pink lines). Panel 2: data from panel 1 were normalized to Z-scores and super imposed on the same axes. False discovery rate (FDR) of 0.05 is depicted by a dotted blue line. Panels 3 and 4: 30-, 60-, and 90-minute hydroxyurea synchronized samples were subjected to Illumina whole genome sequencing and the relative read depth (S/G1) was plotted (panel 3: WT gray color gradient and panel 4: Chaos3 pink color gradient).

**Supplemental Figure 3:**
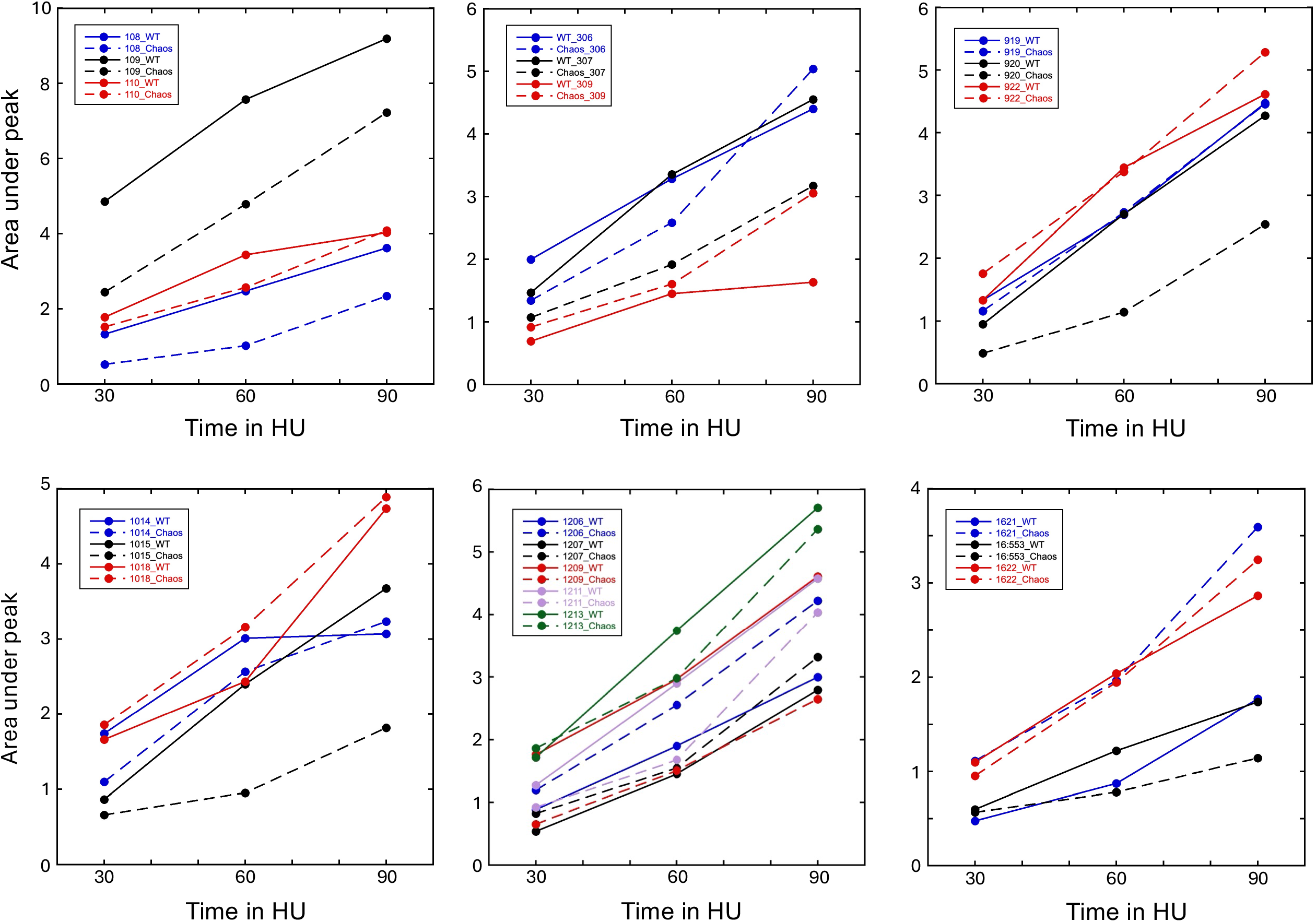
Replication initiation defect of *mcm4^Chaos3^* sensitive origins. (A) Short read sequencing data of a 30-, 60-, and 90-minute (x-axis) hydroxyurea, synchronized culture (Figure 2, Supplemental Fig 2) was used to determine the area under the peak value (y-axis) for the centromere-proximal, *mcm4^Chaos3^* sensitive origins and their flanking origins plotted as colored circles with either solid (WT) or dotted lines (*mcm4^Chaos3^*). A plot was made for each of the identified centromere-proximal, *mcm4^Chaos3^* sensitive origins (chr I, III, IX, X, XII, XVI). See Fig 2B for Chr V.

**Supplemental Figure 4:**
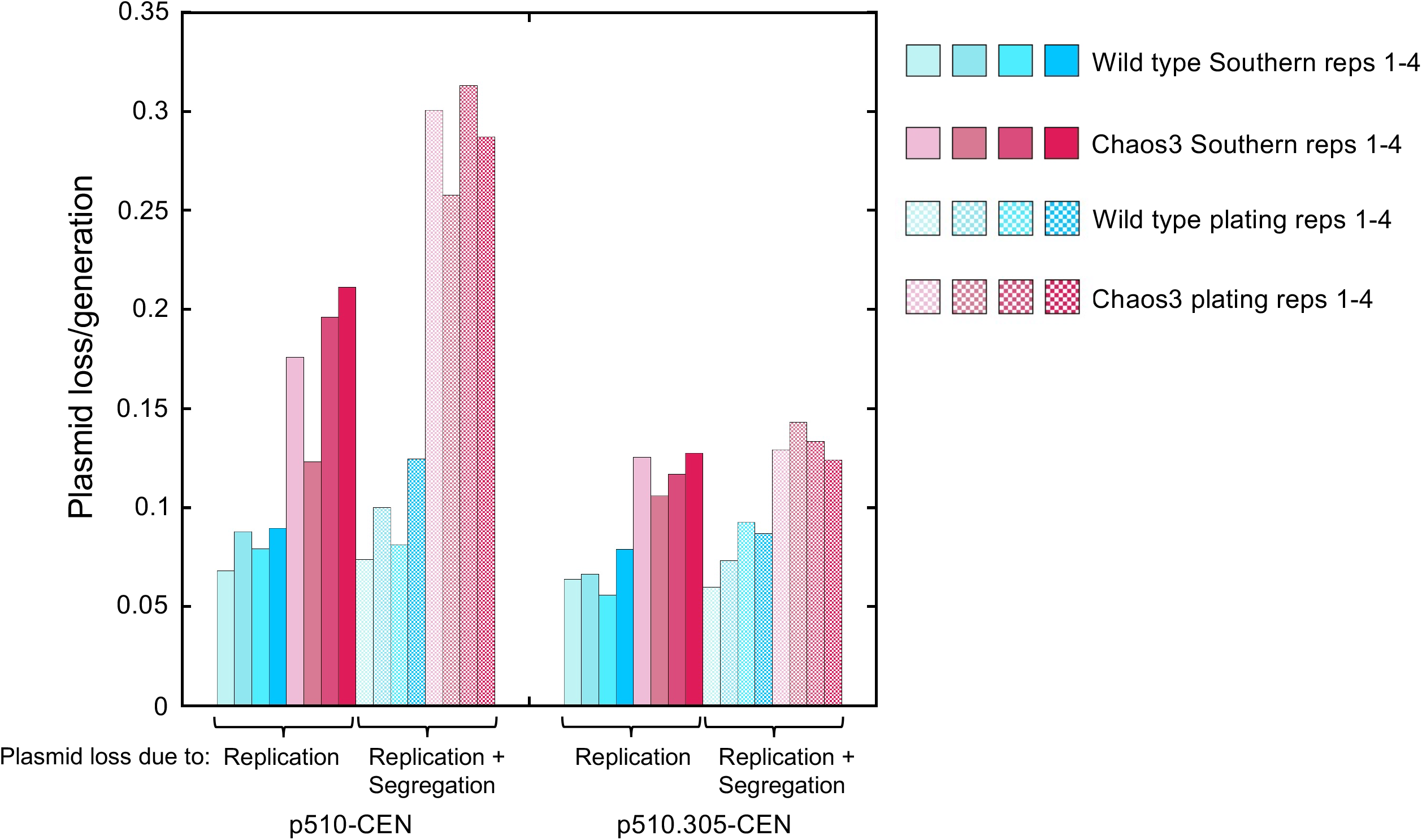
Individual replicates demonstrating variation in plasmid maintenance. (C) *ARS510* (p510-CEN) and *ars510Δ::ARS305* (p510.305-CEN) plasmid loss rates for replicates 1-4 distinguishes plasmid loss due to replication and segregation errors (wild type, dotted blue color gradient and Chaos3, dotted pink color gradient) from replication errors (wild type, solid blue color gradient and Chaos3, solid pink color gradient).

**Supplemental Figure 5:**
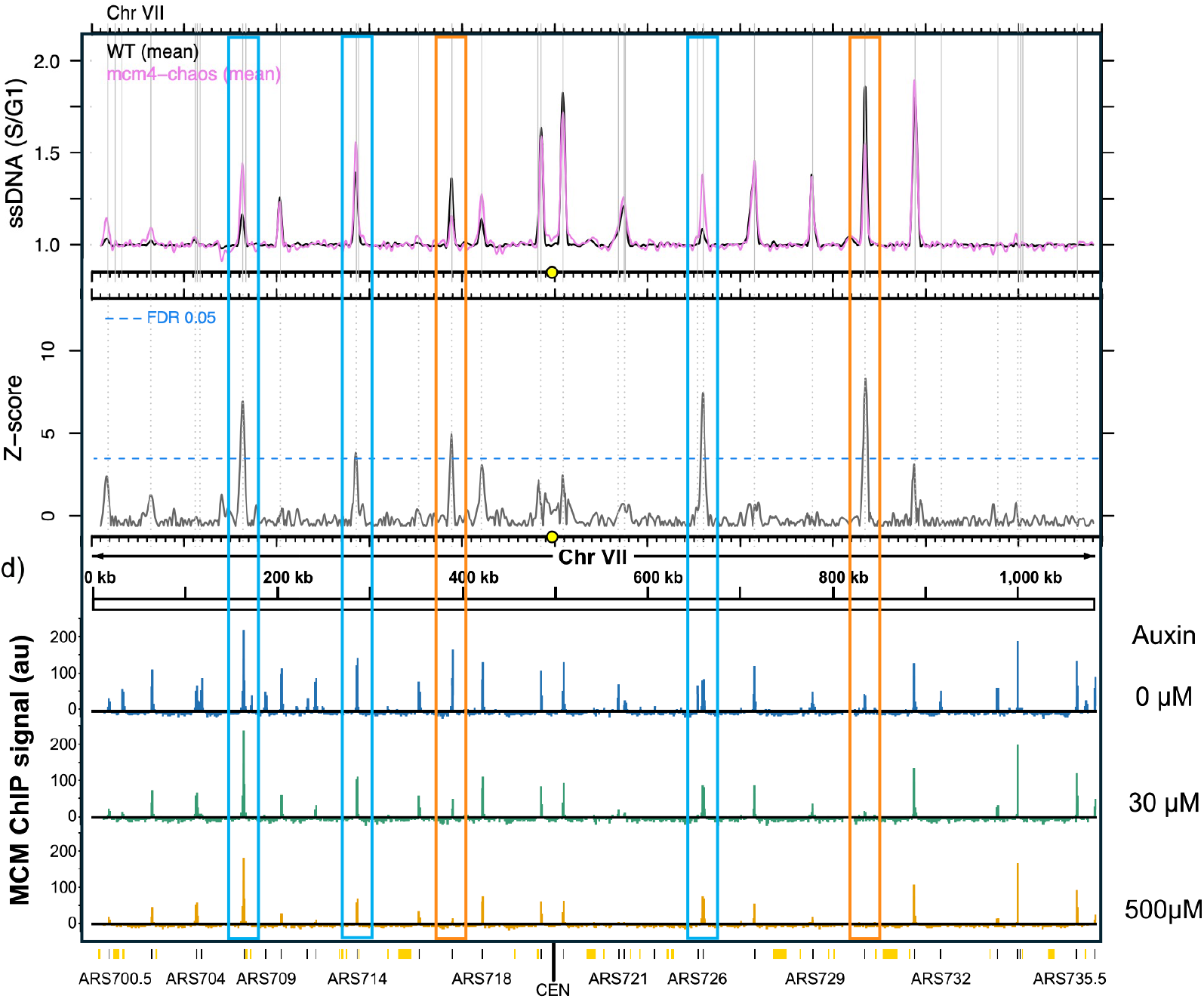
Comparison of early S-phase origin firing to MCM depletion data. (A) Comparison of origin firing (top panel, ssDNA) and Z-scores (middle panel) to Dukaj and Rhind’s MCM depletion data (bottom panel) (43). For the MCM depletion MNase-ChIP-seq coverage on Chromosome VII was normalized in 1 kb bins. ARS-annotated origins are shown as black lines and potential, non-experimentally confirmed origins are shown as yellow lines/boxes. Blue boxes highlight origins that have a significant increase in firing based on ssDNA and Z-score and maintain MCM ChIP signal. Orange boxes highlight origins that have a significant decrease in firing based on ssDNA and Z-score and have diminished MCM ChIP signal. MCM depletion reproduced from Dukaj and Rhind (43) under CCBY 4.0 license (https://creativecommons.org/licenses/by/4.0/).

## Supplemental files

Supplemental file 1: List of strains used

Supplemental file 2: List of primers used

Supplemental file 3: Raw data used to generate all figures

