## Supplemental file 1 for "Making sense of chaos: replication and segregation defects in the yeast *mcm4^Chaos^*^3^ mutant"

| Strain Name | Genotype (All strains derived from A364a) | Source |
| --- | --- | --- |
| HM14-3a | MAT**a**, CDC7, URA3, CAN1S, bar1, trp1-289, leu2-3,112, his6, MCM4 | Heather McCune |
| C7* | MAT**a**, CDC7, ura3-52, can1-100, bar1, trp1-289, leu2-3,112, his6, *mcm4^Chaos3^* | Batool Mutawe |
| AKM001 | MAT**ɑ**, CDC7, ura3-52, can1-100, bar1, trp1-289, leu2-3,112, his6, MCM4 | This study |
| AKM007 | MAT**a**, CDC7, URA3, NAT*CAN1S, ubp9Δ::CAN1S, bar1, trp1-289, leu2-3,112, his6, MCM4 | This study |
| AKM015 | MAT**ɑ**, CDC7, ura3-52, can1-100, bar1, trp1-289, leu2-3,112, his6, *mcm4^Chaos3^* | This study |
| AKM019 | MAT**a**, CDC7, URA3, NAT*CAN1S, ubp9Δ::CAN1S, bar1, trp1-289, leu2-3,112, his6, MCM4, ctf19Δ::KanMx | This study |
| AKM020 | MAT**a**, CDC7, URA3, NAT*CAN1S, ubp9Δ::CAN1S, bar1, trp1-289, leu2-3,112, his6, MCM4, ctf19Δ::KanMx | This study |
| AKM021 | MAT**ɑ**, CDC7, ura3-52, can1-100, bar1, trp1-289, leu2-3,112, his6, MCM4, ctf19Δ::KanMx | This study |
| AKM022 | MAT**a**, CDC7, ura3-52, can1-100, bar1, trp1-289, leu2-3,112, his6, MCM4, ctf19Δ::KanMx | This study |
| AKM023 | MAT**ɑ**, CDC7, ura3-52, can1-100, bar1, trp1-289, leu2-3,112, his6, MCM4, ctf19Δ::KanMx | This study |
| AKM025 | MAT**a**, CDC7, URA3, NAT*CAN1S, ubp9Δ::CAN1S, bar1, trp1-289, leu2-3,112, his6, *mcm4^Chaos3^* | This study |
| AKM026 | MAT**ɑ**, CDC7, ura3-52, can1-100, bar1, trp1-289, leu2-3,112, his6, *mcm4^Chaos3^*, ctf19Δ::KanMx | This study |
| AKM032 | MAT**a**, CDC7, URA3, NAT*CAN1S, ubp9Δ::CAN1S, bar1, trp1-289, leu2-3,112, his6, *mcm4^Chaos3^*, ctf19Δ::KanMx | This study |
| AKM037 | MAT**a**, CDC7, ura3-52, can1-100, bar1, trp1-289, leu2-3,112, his6, *mcm4^Chaos3^*, ctf19Δ::KanMx | This study |
| AKM044 | MAT**a**, CDC7, URA3, NAT*CAN1S, ubp9Δ::CAN1S, bar1, trp1-289, leu2-3,112, his6, MCM4, *ars510Δ::ARS305* | This study |
| AKM048 | MAT**a**, CDC7, URA3, NAT*CAN1S, ubp9Δ::CAN1S, bar1, trp1-289, leu2-3,112, his6, *mcm4^Chaos3^*, *ars510Δ::ARS305* | This study |
| JIV005 | MAT**a**, CDC7, URA3, NAT*CAN1S, ubp9Δ::CAN1S, bar1, trp1-289, leu2-3,112, his6, MCM4, *ars510Δ::ARS305*, ctf19Δ::KanMx | This study |
| JIV006 | MAT**a**, CDC7, URA3, NAT*CAN1S, ubp9Δ::CAN1S, bar1, trp1-289, leu2-3,112, his6, *mcm4^Chaos3^*, *ars510Δ::ARS305*, ctf19Δ::KanMx | This study |
| JIV008 | MAT**a**, CDC7, URA3, NAT*CAN1S, ubp9Δ::CAN1S, bar1, trp1-289, leu2-3,112, his6, MCM4, *ars510Δ::ARS305^fkh inv^* | This study |
| JIV021 | MAT**a**, CDC7, URA3, NAT*CAN1S, ubp9Δ::CAN1S, bar1, trp1-289, leu2-3,112, his6, *mcm4^Chaos3^*, *ars510Δ::ARS305^fkh inv^* | This study |
| JIV036 | MAT**a**, CDC7, ura3-52, can1-100, bar1, trp1-289, leu2-3,112, his6, MCM4 | This study |
| AD24-27d | MAT**ɑ**, CDC7, URA3, CAN1S, bar1, trp1-289, leu2-3,112, his6, MCM4 | This study |
| MR98-2b | MAT**ɑ**, CDC7, URA3, CAN1S, bar1, trp1-289, leu2-3,112, his6, *mcm4^Chaos3^* | This study |
| AMD003 | MAT**a**/MAT**ɑ**, CDC7/CDC7, URA3/ura3-52, NAT*CAN1S/can1-100, ubp9Δ::CAN1S/UBP9, bar1/bar1, trp1-289/trp1-289, leu2-3,112/leu2-3,112, his6/his6, MCM4/MCM4 | This study |
| AMD019 | MAT**a**/MAT**ɑ**, CDC7/CDC7, URA3/ura3-52, NAT*CAN1S/can1-100, ubp9Δ::CAN1S/UBP9, bar1/bar1, trp1-289/trp1-289, leu2-3,112/leu2-3,112, his6/his6, *mcm4^Chaos3^*/ *mcm4^Chaos3^* | This study |
| AMD020 | MAT**a**/MAT**ɑ**, CDC7/CDC7, URA3/ura3-52, NAT*CAN1S/can1-100, ubp9Δ::CAN1S/UBP9, bar1/bar1, trp1-289/trp1-289, leu2-3,112/leu2-3,112, his6/his6, MCM4/MCM4, ctf19Δ::KanMx/ ctf19Δ::KanMx | This study |
| AMD033 | MAT**a**/MAT**ɑ**, CDC7/CDC7, URA3/ura3-52, NAT*CAN1S/can1-100, ubp9Δ::CAN1S/UBP9, bar1/bar1, trp1-289/trp1-289, leu2-3,112/leu2-3,112, his6/his6, *mcm4^Chaos3^*/ *mcm4^Chaos3^*, ctf19Δ::KanMx/ ctf19Δ::KanMx | This study |
| AMD040 | MAT**a**/MAT**ɑ**, CDC7/CDC7, URA3/ura3-52, NAT*CAN1S/can1-100, ubp9Δ::CAN1S/UBP9, bar1/bar1, trp1-289/trp1-289, leu2-3,112/leu2-3,112, his6/his6, MCM4/MCM4, *ars510Δ::ARS305*/*ARS510* | This study |
| JVD009 | MAT**a**/MAT**ɑ**, CDC7/CDC7, URA3/ura3-52, NAT*CAN1S/can1-100, ubp9Δ::CAN1S/UBP9, bar1/bar1, trp1-289/trp1-289, leu2-3,112/leu2-3,112, his6/his6, MCM4/MCM4, *ars510Δ::ARS305*/*ARS510*, ctf19Δ::KanMx/ ctf19Δ::KanMx | This study |
| JVD028 | MAT**a**/MAT**ɑ**, CDC7/CDC7, URA3/ura3-52, NAT*CAN1S/can1-100, ubp9Δ::CAN1S/UBP9, bar1/bar1, trp1-289/trp1-289, leu2-3,112/leu2-3,112, his6/his6, *mcm4^Chaos3^*/ *mcm4^Chaos3^*, *ars510Δ::ARS305*/*ARS510*, ctf19Δ::KanMx/ ctf19Δ::KanMx | This study |
| JVD031 | MAT**a**/MAT**ɑ**, CDC7/CDC7, URA3/ura3-52, NAT*CAN1S/can1-100, ubp9Δ::CAN1S/UBP9, bar1/bar1, trp1-289/trp1-289, leu2-3,112/leu2-3,112, his6/his6, *mcm4^Chaos3^*/ *mcm4^Chaos3^*, *ars510Δ::ARS305^fkh inv^*/*ARS510* | This study |
| JVD037 | MAT**a**/MAT**ɑ**, CDC7/CDC7, URA3/ura3-52, NAT*CAN1S/can1-100, ubp9Δ::CAN1S/UBP9, bar1/bar1, trp1-289/trp1-289, leu2-3,112/leu2-3,112, his6/his6, *mcm4^Chaos3^*/ *mcm4^Chaos3^*, *ars510Δ::ARS305*/*ARS510* | This study |
| JVD041 | MAT**a**/MAT**ɑ**, CDC7/CDC7, URA3/ura3-52, NAT*CAN1S/can1-100, ubp9Δ::CAN1S/UBP9, bar1/bar1, trp1-289/trp1-289, leu2-3,112/leu2-3,112, his6/his6, MCM4/MCM4, *ars510Δ::ARS305^fkh inv^*/*ARS510* | This study |
