## Supplemental file 2 for "Making sense of chaos: replication and segregation defects in the yeast *mcm4^Chaos^*^3^ mutant"

| **Primer number** | **Sequence 5’ —> 3’** | **Usage** |
| --- | --- | --- |
| 1 | CAGTCTTTATATCAGTCTTTGCATGGCTTTGCATCTGATGCTGGCTCTACAGATTGTACTGAGAGTGCAC | Amplification of CloNAT from pFvL99 |
| 2 | TGGATAGTATATACGTCAAATTCTTTTGGTATTTTATCGCCGAGAAGTCGctgtgcggtatttcacaccg |  |
| 3 | AGAAACACAGGCTGATA | Amplification of CAN1^S^ |
| 4 | GGTGCTAAGGAGGTTAT |  |
| 5 | GATCAGGAACAATAGAAAGAAACGGTTTTAGAGCTAG | CRISPR gRNA targeting UBP9 for CAN1^S^ insertion |
| 6 | CTAGCTCTAAAACCGTTTCTTTCTATTGTTCCT |  |
| 7 | ATATGGATTTTACGTCTGATACGACGAATTCGCACGACACagattgtactgagagtgcac | Amplification of KanMX from pRS400 |
| 8 | TCACCTGGCGTACATGTCCGGGAATAGGCAAACGTTGCAGctgtgcggtatttcacaccg |  |
| 9 | GGCAAGAGGGCTTCGCCGGCTTTTACTCTGGGTTTGCAACTAATTTGGTAAGGACAGTACCTGCTGCTGTGGTAACACTATATCGTGTAAGCTGGGGTGA | Amplification of ARS305 with homology ends to ARS510 |
| 10 | ACACCAAATGTTCTTTAAAAAACTCAATATCTTTGACTTAAAATTTAACTTACATGTTAATGGTATGTAAGATATACTTGGTAGCACTTTGATGAGGTCT |  |
| 11 | GATCCTTGTTCGAATAGTAAAAGAGTTTTAGAGCTAG | CRISPR gRNA targeting ARS510 for ARS305 replacement |
| 12 | CTAGCTCTAAAACTCTTTTACTATTCGAACAAG |  |
| 13 | TTCAAAAGAAATTACTCAAATATGTCCAGTTTCATGTACTGTCCGATGTGATTTATTATTTTTGTAAATATTTGTAGTTCTTAAAGCTAAG | ARS305 fkh inv template |
| 14 | TCTTTGGAGCTCAAGTGGATTGAGGCCACAGCAAGACCGAGTTTGAATGCTCAACTCTTCAAAAGAAATTACTCAAA | Extended ARS305 fkh inv template for higher transformation efficiency |
| 15 | CCCTTTAATTTTAGGATATGAAAACAAGAATTTATCAAAGAAAAAAATCTTAGCTTTTACAAA |  |
| 16 | GATCGTCCAGTTTCATGTACTGTCGTTTTAGAGCTAG | CRISPR gRNA targeting ars510Δ::ARS305 for ARS305 fkh inversion |
| 17 | CTAGCTCTAAAACGACAGTACATGAAACTGGAC |  |
| 18 | CCTCGTCAATGTCCCTACTTAC | 2D gel and plasmid loss Southern blot probe |
| 19 | CAACAACCTCCGAACACTAGA |  |
| 20 | GGGTACCAACGTCAGTATCC | Co-culture PCR for sequencing |
| 21 | GTGCCGCCAAATAGCTGAAG |  |
